# Impacts of yeast Tma20/MCTS1, Tma22/DENR and Tma64/eIF2D on translation reinitiation and ribosome recycling

**DOI:** 10.1101/2024.03.06.583729

**Authors:** Kristína Jendruchová, Swati Gaikwad, Kristýna Poncová, Stanislava Gunišová, Leoš Shivaya Valášek, Alan G. Hinnebusch

## Abstract

Recycling of 40S ribosomal subunits following translation termination, entailing release of deacylated tRNA and dissociation of the empty 40S subunit from mRNA, involves yeast Tma20/Tma22 heterodimer and Tma64, counterparts of mammalian MCTS1/DENR and eIF2D. MCTS1/DENR enhance reinitiation at short upstream open reading frames (uORFs) harboring penultimate codons that confer dependence on these factors in bulk 40S recycling. Tma factors, by contrast, inhibited reinitiation at particular uORFs in extracts; however, their roles at regulatory uORFs in vivo were unknown. We examined effects of eliminating Tma proteins on reinitiation at regulatory uORFs mediating translational control of *GCN4* optimized for either promoting (uORF1) or preventing (uORF4) reinitiation. We found that the Tma proteins generally impede reinitiation at native uORF4 and uORF4 variants equipped with various penultimate codons regardless of their Tma-dependence in bulk recycling. The Tma factors have no effect on reinitiation at native uORF1, and equipping uORF1 with Tma-dependent penultimate codons generally did not confer Tma-dependent reinitiation; nor did converting the uORFs to AUG-stop elements. Thus, effects of the Tma proteins vary depending on the reinitiation potential of the uORF and the penultimate codon, but unlike in mammals, are not principally dictated by the Tma-dependence of the codon in bulk 40S recycling.

## INTRODUCTION

Synthesis of proteins is carried out by ribosomes, molecular machines that assemble from the small (40S) and large (60S) ribosomal subunits. During translation initiation, the 40S subunit acquires the ternary complex (TC) composed of the methionyl initiator Met-tRNAi^Met^, GTP and the eukaryotic Initiation Factor (eIF) 2, forming the 43S pre-initiation complex (PIC). Upon mRNA recruitment, the 48S PIC thus formed scans the mRNA 5’ leader region to locate the authentic initiation codon, generally an AUG triplet in a favorable “Kozak” sequence context. Subsequently, subunit joining occurs to produce the 80S initiation complex and elongation commences. Upon stop codon recognition, translation terminates, and the nascent peptide is released. This is followed by ribosomal recycling, i.e., splitting of both ribosomal subunits and dissociation of the 40S and the deacylated tRNA that decoded the last sense codon from mRNA, to start the new translational cycle (reviewed in^1,2,3,4^). In some cases, the recycling step is incomplete, leaving the post-termination 40S mRNA-bound, which allows it to resume traversing downstream and, upon reacquisition of the TC, to initiate translation of a downstream ORF in a process called reinitiation (REI) (reviewed in^5^).

Translation termination and ribosome recycling are intrinsically linked. When the stop codon enters the ribosomal A site, it is recognized by a heterodimer composed of eukaryotic Release Factors 1 and 3 (eRF1/eRF3). Stop codon recognition results in GTP hydrolysis on eRF3 followed by its release. eRF1 then catalyzes hydrolysis of the nascent polypeptide chain, which generates the post-termination 80S ribosome harboring deacylated tRNA base-paired with the penultimate codon in the P site^6^. The ATP-binding cassette protein Rli1/ABCE1 (yeast/mammals) facilitates the dissociation of the 60S subunit both *in vitro* in yeast and mammalian reconstituted systems^7,8^ and *in vivo* in yeast cells^9^, aided by the eIF3-associated factor eIF3j^10,11^. Studies *in vitro* revealed that after subunit splitting, the deacylated tRNA and mRNA are dissociated from the 40S post-termination complex by the non-canonical initiation factor eIF2D or its functionally and structurally related heterodimer MCTS1/DENR^12,13^. DENR and MCTS1 bear sequence similarity to the N-terminal and C-terminal regions of eIF2D, respectively. The MCTS1 and the eIF2D N-terminal region harbor DUF1974 and PUA domains, whereas DENR and eIF2D’s C-terminal region contain SWIB/MDM2 and SUI/eIF1 domains. eIF2D additionally contains a central winged-helix domain^4^.

Ribosome profiling studies demonstrated that the yeast orthologs of eIF2D, MCTS1 and DENR, known as Tma64, Tma20 and Tma22, respectively^14^, function in 40S post-termination complex recycling *in vivo*^15,16^. Thus, deleting the corresponding genes led to accumulation of unrecycled 40S subunits at the majority of stop codons, with the largest defect observed in *Δtma20Δtma64* (*tmaΔΔ*) cells. Quantifying the accumulation of unrecycled 40S subunits at all individual stop codons in *tmaΔΔ* versus WT cells revealed that certain penultimate codons result in significantly higher accumulation of unrecycled 40S subunits in the mutant cells—a phenomenon dubbed Tma-dependence. Furthermore, comparison of the double deletant to single deletant strains showed that Tma64 is largely dispensable, whereas both subunits of the Tma20/Tma22 heterodimer are required for the majority of 40S recycling events *in vivo*. A notable consequence of defective 40S recycling in mutants lacking an intact Tma20/Tma22 heterodimer is increased REI downstream of stop codons of main ORFs within the 3’ untranslated regions (3’ UTRs) of mRNAs. Such non-canonical REI frequently occurs by conventional scanning of the unrecycled post-termination 40S complexes to 3’ UTR-situated AUG codons, preferentially in optimum Kozak context^15,16^. Absence of the Tma20/Tma22 heterodimer also conferred increased canonical REI downstream of the stop codons of upstream open reading frames (uORFs) in the 5’ UTRs of reporter mRNAs in yeast cell-free translation extracts^15^.

Whereas yeast Tma20/Tma22 were shown to inhibit REI, work on their mammalian orthologs indicated that human DENR conversely promotes REI after translation of certain uORFs, including 1aa-long uORFs comprised of only a start and stop codon, referred to here as ‘start-stops’, with their ATG start codons in optimum Kozak context^17,18^. In the absence of DENR/MCTS1, mRNAs containing start-stops exhibit reduced translation of the downstream main ORF. Based on 40S ribosome profiling data, DENR-dependence for REI was also established for certain other short uORFs in a manner generally dictated by the presence of particular penultimate codons that confer increased dependence on DENR for bulk 40S recycling in the translatome, the efficiency of which in turn determined the efficiency of REI^13^. Indeed, the DENR-dependence often inversely correlates with the propensity of the deacylated tRNA decoding the penultimate codon to dissociate spontaneously from post-termination 40S complexes in reconstituted termination/recycling systems^13,19^. Accordingly, it was proposed that uORFs harboring penultimate codons with low rates of spontaneous dissociation of the cognate tRNAs require DENR/MCTS1 to accelerate release of the deacylated tRNA to enable the post-termination 40S to resume traversing downstream and reacquire the TC at the now-empty P site, enabling REI. The DENR-requirement for REI included the regulatory short uORF that stimulates reinitiation on *ATF4* mRNA (uORF1), allowing scanning ribosomes to bypass a more distally positioned uORF (uORF2) that inhibits initiation at the main CDS, both in mammalian cells^13^ and fruit flies^20^.

To reconcile the apparent discrepancy between yeast Tma factors and their mammalian counterparts, it has been suggested^13^ that release of the deacylated tRNA by MCTS1/DENR does not lead to dissociation of post-termination 40S subunits at relatively short uORFs in mammalian cells owing to retention of eIF3, eIF4G1, or eIF4E by 80S ribosomes translating the uORFs and continued occupancy of these factors on the post-termination 40S subunits^21,22,23^. These retained eIFs presumably impede the 40S dissociation function of MCTS1/DENR, making the only observable consequence of depleting DENR a reduction in REI at the subset of uORFs requiring DENR for release of the deacylated tRNA, whose continued presence prevents the 40S post-termination complexes from traversing downstream or recruiting TC to the P site. Depleting DENR has no effect on REI at the short uORFs where deacylated tRNA dissociates independently of MCTS1/DENR. These expected outcomes are depicted schematically in panels (i)(a) and (ii)(a) of Fig. 1A (cf. (+) DENR/Tma and (-) DENR/Tma). In yeast, by contrast, release of the deacylated tRNA, whether or not it is dependent on the Tma proteins, would lead to subsequent dissociation of post-termination 40S subunits from mRNA and low-level REI following most uORFs in WT cells, because eIFs are generally not retained by 80S ribosomes translating yeast uORFs^24,25,26,27^. As such, a further reduction in REI in cells lacking Tma factors is not expected for typical yeast uORFs even when they depend on the Tma factors to stimulate release of the deacylated tRNAs from the post-termination 40S subunits. These expected outcomes for typical uORFs in yeast are shown in panels (i)(a) and (ii)(a) of Fig. 1B (cf. WT and *tmaΔΔ*).

**FIGURE 1:**
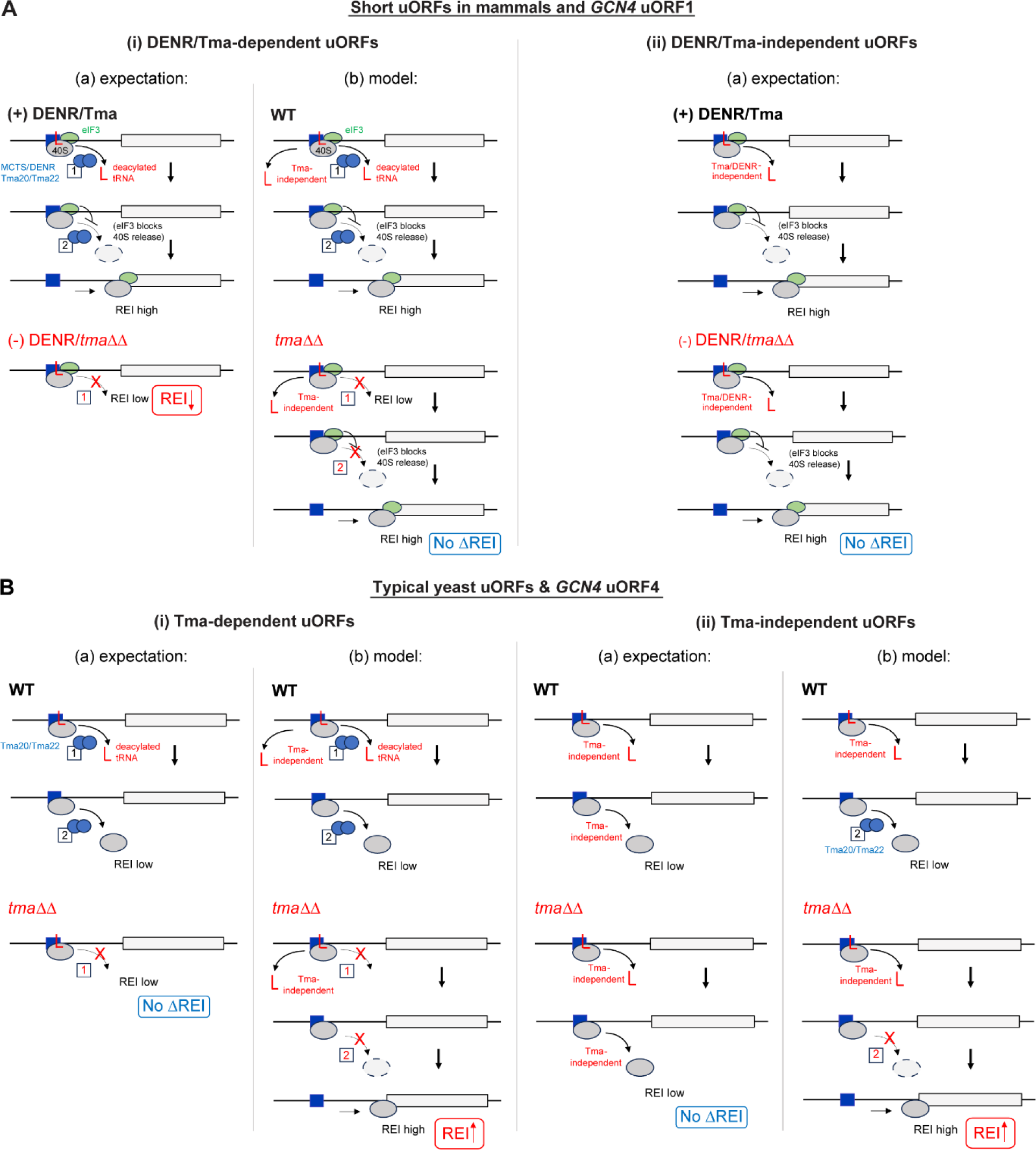
Proposed mechanisms governing MCTS1/DENR mediated recycling and REI at mammalian short uORFs and expected and proposed functions for yeast Tma20/Tma22 at REI-permissive *GCN4* uORF1 and REI-nonpermissive *GCN4* uORF4. MCTS1/DENR or Tma20/Tma22 are depicted as linked blue ovals whose distinct functions in releasing the penultimate deacylated tRNA (red “L”) and subsequently dissociating the empty 40S from the mRNA are labeled with boxes 1 and 2, respectively, and solid or dashed arrows. **A)** DENR/Tma-mediated recycling at short mammalian uORFs and *GCN4* uORF1 depending on the presence of (i) DENR/Tma-dependent penultimate codons requiring the recycling factors for tRNA release, or (ii) DENR/Tma-independent codons where tRNA can be released spontaneously. **(i) DENR/Tma-dependent uORFs. (a) expectation**: WT recycling factors in “(+) DENR/Tma” cells release the penultimate tRNA but the second function in 40S dissociation is impeded by eIF3 (green oval) retained on post-termination 40S subunits to enable high-level REI at the main CDS. In “(-) DENR/*tmaΔΔ*” cells lacking the factors, failure to release the penultimate tRNA lowers REI. **(b) model:** Based on findings here we propose that deacylated tRNA can be released from uORF1 variants by both Tma-dependent and-independent mechanisms in WT yeast, and this Tma-independent tRNA release maintains high-level REI in *tmaΔΔ* cells, as 40S dissociation is not impeded at uORF1 in neither WT nor mutant cells. **(ii) DENR/Tma-independent uORFs. (a) expectation:** release of the tRNA and high-level REI occurs in the presence (top) or absence (bottom) of the recycling factors, yielding no decrease in REI in *tmaΔΔ* vs. WT cells. **B)** Tma-mediated recycling at typical yeast short uORFs and *GCN4* uORF4 depending on the presence of (i) Tma-dependent penultimate codons, or (ii) Tma-independent codons. **(i) Tma-dependent uORFs. (a) expectation**: In WT cells, Tma factors release the penultimate tRNA but also dissociate the empty 40S subunit from mRNA, leading to low-level REI. In *tmaΔΔ* cells, the absence of Tma-mediated tRNA release helps to ensure low-level REI, yielding no change in REI in *tmaΔΔ* vs. WT cells. **(b) model:** Based on our findings we propose that tRNA can be released from uORF4 variants by Tma-dependent and-independent mechanisms in WT, and Tma-independent tRNA release enables increased REI in *tmaΔΔ* cells where the 40S dissociation function is absent. **(i) Tma-independent uORFs. (a) expectation:** Tma-independent tRNA release occurs in both WT and *tmaΔΔ* cells, but REI is low in both cases because the empty 40S subunits also dissociate from the mRNA independently of Tma factors, for no change in REI vs. WT. **(b) model:** Our findings suggest that in WT cells 40S dissociation is catalyzed by Tma factors even at such Tma-independent uORFs, and that absence of this function in *tmaΔΔ* cells, coupled with Tma-independent tRNA release, enables increased REI in the mutant.

A different outcome would be predicted for atypical uORFs in yeast where 40S dissociation is counteracted by initiation factors that remain associated with post-termination 40S complexes. The 5’-proximal AUG-initiated uORF in *GCN4* mRNA, uORF1, is the best characterized such uORF in yeast, being optimized for eIF3 and eIF4G binding and retention of post-termination 40S subunits by the presence of cis-acting reinitiation promoting elements (RPEs) rendering it highly permissive for REI downstream (Fig. 2)^5,27,28^. Accordingly, the 40S post-termination complexes at the uORF1 stop codon should not spontaneously dissociate from mRNA on release of the deacylated tRNA, enabling high-level REI in WT cells whether or not tRNA release depends on the Tma factors (Fig. 1A(i)(a)-(ii)(a), (+) DENR/Tma). This will allow us to determine whether eliminating the Tma proteins confers reduced REI at uORF1 variants equipped with penultimate triplets requiring Tma factors for release of deacylated tRNA, while having no effect at uORF1 variants containing Tma-independent triplets (Fig. 1A(i)(a)-(ii)(a), (-) DENR/*tmaΔ*). Indeed, *GCN4* uORF1 is functionally equivalent to uORF1 in *ATF4* mRNA, which was shown to be DENR-dependent for efficient REI^13,20^; and *GCN4* and *ATF4* translation are governed by very similar REI mechanisms^5^. So far, we have determined that the Tma20/Tma22 heterodimer is unlikely to be required to dissociate the deacylated tRNA^Cys^ from the native penultimate UGC codon of *GCN4* uORF1, as translational control of WT *GCN4* mRNA occurred normally in the *tma20Δtma64Δ* mutant^29^. Consistently, UGC was judged to be DENR-independent in mammalian cells^13^ and to be one of the least Tma-dependent codons for yeast 40S recycling^16^. In addition, there is evidence that the cognate tRNA^Cys^ is weakly associated with mammalian 40S post-termination complexes^19^. Here, we examine whether substituting the native UGC codon in uORF1 with other codons judged to be Tma-dependent for bulk 40S recycling reduces REI in *tmaΔΔ* cells, as observed for DENR-dependent short uORFs in DENR-depleted mammalian cells (Fig. 1A(i)(a)).

**FIGURE 2:**
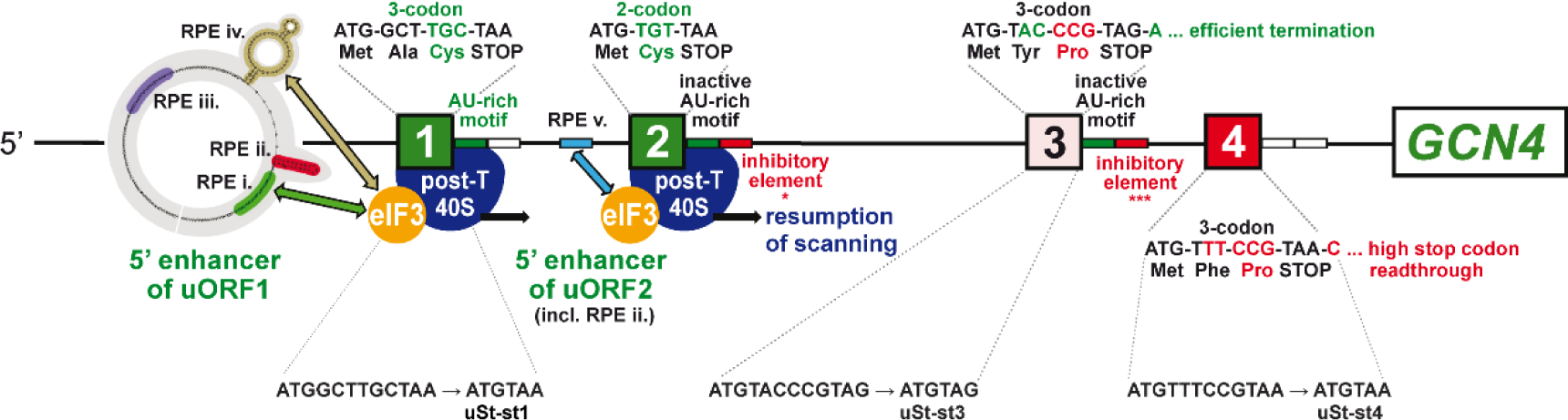
Summary of all *cis*-determinants that either promote or inhibit reinitiation on *GCN4* mRNA after translation of its four short uORFs. Schematics of the 5′ enhancers of uORF1 and 2 containing their respective RPEs, some of which functionally interact with eIF3 to promote resumption of scanning. Green color-coding generally indicates stimulatory effects of the corresponding *cis*-factors on efficiency of REI, whereas red color-coding indicates inhibitory effects (with the exception of RPE ii. of uORF1, which is also stimulatory); the number of asterisks below the inhibitory elements of the uORF2 and uORF3 3′ sequences depicts the degree of their inhibition as determined experimentally. Mutations converting uORFs 1, 3, or 4 to Start-stop elements uSt-st1, uSt-st3, or uSt-st4, described later in RESULTS, are given below the respective WT uORF schematics. Reprinted and modified with permission from^33^.

In contrast to REI-permissive uORF1, the fourth AUG-initiated uORF in *GCN4* mRNA (uORF4) does not allow retention of eIFs on post-termination 40S subunits, owing to multiple, surrounding cis-acting sequences, thus making it non-permissive for REI and repressive to *GCN4* translation in WT cells^26,27,30,31,32,33^. We reasoned that uORF4 is typical of short uORFs in yeast that do not allow REI owing to efficient dissociation of 40S post-termination complexes regardless of the requirement for Tma factors in releasing tRNA (Fig.1B(i)(a)-(ii)(a), WT). uORF4 contains a penultimate CCG proline codon, whose dependence on DENR for recycling and REI in human cells is ambiguous^13^, and was neither unusually dependent nor independent of the Tma proteins (i.e. Tma-neutral) for bulk 40S recycling in yeast^16^. Since elimination of the Tma20/Tma22 heterodimer did not impact *GCN4* translational control^29^, it seems unlikely that CCG contributes to low-level REI at WT uORF4 by imposing a requirement for Tma factors for tRNA release. Here, we have tested the expectation that replacing the native CCG codon with other codons known to be Tma-dependent for bulk recycling will not reduce REI in *tmaΔΔ* cells because, lacking eIF3, the 40S subunits will dissociate and fail to reinitiate regardless of tRNA release (Fig. 1B(i)(a), *tmaΔΔ* versus WT). We have also examined whether converting the native 3-codon uORF1 and uORF4 to start-stop elements by eliminating their second and third coding triplets confers a dependence on the Tma proteins for REI in the manner observed for DENR with these specialized uORFs in mammalian cells^17,18^.

Our results depart from the expected outcomes for *GCN4* uORF4 shown in Fig. 1B and suggest a role for Tma factors at this uORF in dissociating 40S post-termination complexes from mRNA following release of deacylated tRNA, regardless of whether the penultimate triplets are Tma-dependent in bulk recycling. Our findings also differ from those obtained for short, REI-permissive uORFs in mammalian cells^13^ in revealing that *GCN4* uORF1 generally allows efficient REI independently of Tma factors even when equipped with Tma-dependent penultimate codons. We found that REI following start-stop uORFs is also Tma-independent, and we show that Tma64 plays a supportive role to that of Tma20/Tma22 in impeding REI at uORF4, consistent with its auxiliary role in recycling at most stop codons in yeast.

## RESULTS

### Tma64 partially substitutes for the Tma20/Tma22 heterodimer in suppressing REI after translation of the *GCN4* uORF4 variant containing the Tma-dependent TTG^Leu^ codon

To examine the effects of replacing the penultimate codons of uORF1 or uORF4 with Tma-dependent versus Tma-independent triplets, we employed the well-established *GCN4-lacZ* reporter system, which faithfully recapitulates the functions of the *cis* and *trans* regulatory elements/factors involved in *GCN4* translational control^24,25,30,31,32,33,34,35,36,37^. This reporter contains the entire *GCN4* transcription unit, including the ∼600 nt leader harboring the four uORFs (Fig. 2), with the *lacZ* coding sequences inserted in-frame with the *GCN4* main ORF. To study uORF1, we employed the “uORF1-only” reporter lacking the AUGs ofuORFs 2-4. Because leaky-scanning of the uORF1 AUG codon is extremely low^36^, expression of β-galactosidase from this reporter provides a read-out of REI following termination at uORF1. Similarly, uORF4 variants were examined in the “uORF4-only” reporter lacking the AUGs of uORFs 1-3, which produces low levels of β-galactosidase owing to a lack of leaky-scanning and highly inefficient REI at uORF4^34^. Both uORF1 and uORF4 are present at their normal positions in the leader of these reporters.

We first examined the relative contribution of the Tma20/Tma22 heterodimer and Tma64 to the efficiency of REI following uORF4 translation, which we regard as an exemplar of REI-non-permissive uORFs in budding yeas ^3,13^. To this end, uORF4-only reporters were generated (Fig. 3A) with the native penultimate CCG^Pro^ codon replaced by either the TTG^Leu^ codon or TGG^Trp^ codon, judged to be the most dependent or independent of Tma factors, respectively, for bulk 40S recycling in yeast^16^. Assaying the reporters in the three single deletion strains lacking *TMA20, TMA22,* or *TMA64*, the *tma20Δtma22Δtma64Δ* triple deletion mutant, and the isogenic WT revealed that none of the *tma* mutations produced a significant change in expression of the reporter with the Tma-independent TGG^Trp^ codon (Fig. 3B). This result matches the expected outcome depicted in Fig. 1B(ii)(a). At odds with the expectations in Fig. 1B(i)(a), however, the single deletions of *TMA20* or *TMA22* conferred ≥2-fold derepression of the reporter containing the Tma-dependent TTG^Leu^ codon. Although the single deletion of *TMA64* had no effect, the triple mutant showed even higher expression of the TTG^Leu^ reporter compared to the *tma20Δ* and *tma22Δ* single mutants, at >3-fold above the level in WT (Fig. 3B). One way to explain these unexpected findings on the TTG^Leu^ reporter is to propose that eliminating Tma20/Tma22 function in dissociating post-termination 40S subunits from mRNA allows the retained subunits to traverse the leader and reinitiate downstream at the *GCN4* AUG codon, even though the heterodimer’s function in releasing the deacylated leucyl tRNA is absent in the mutant cells. This in turn implies the occurrence of Tma-independent release of the deacylated tRNA at the TTG^Leu^ codon, allowing increased REI to occur when dissociation of the post-termination 40S subunit is impaired by *tma20Δ* or *tma22Δ* mutations. This proposed model is presented in Fig. 1B(i)(b), depicting both Tma-dependent and - independent mechanisms of tRNA release at a Tma-dependent uORF4 variant. Additional evidence supporting this model is presented in the next section.

**FIGURE 3:**
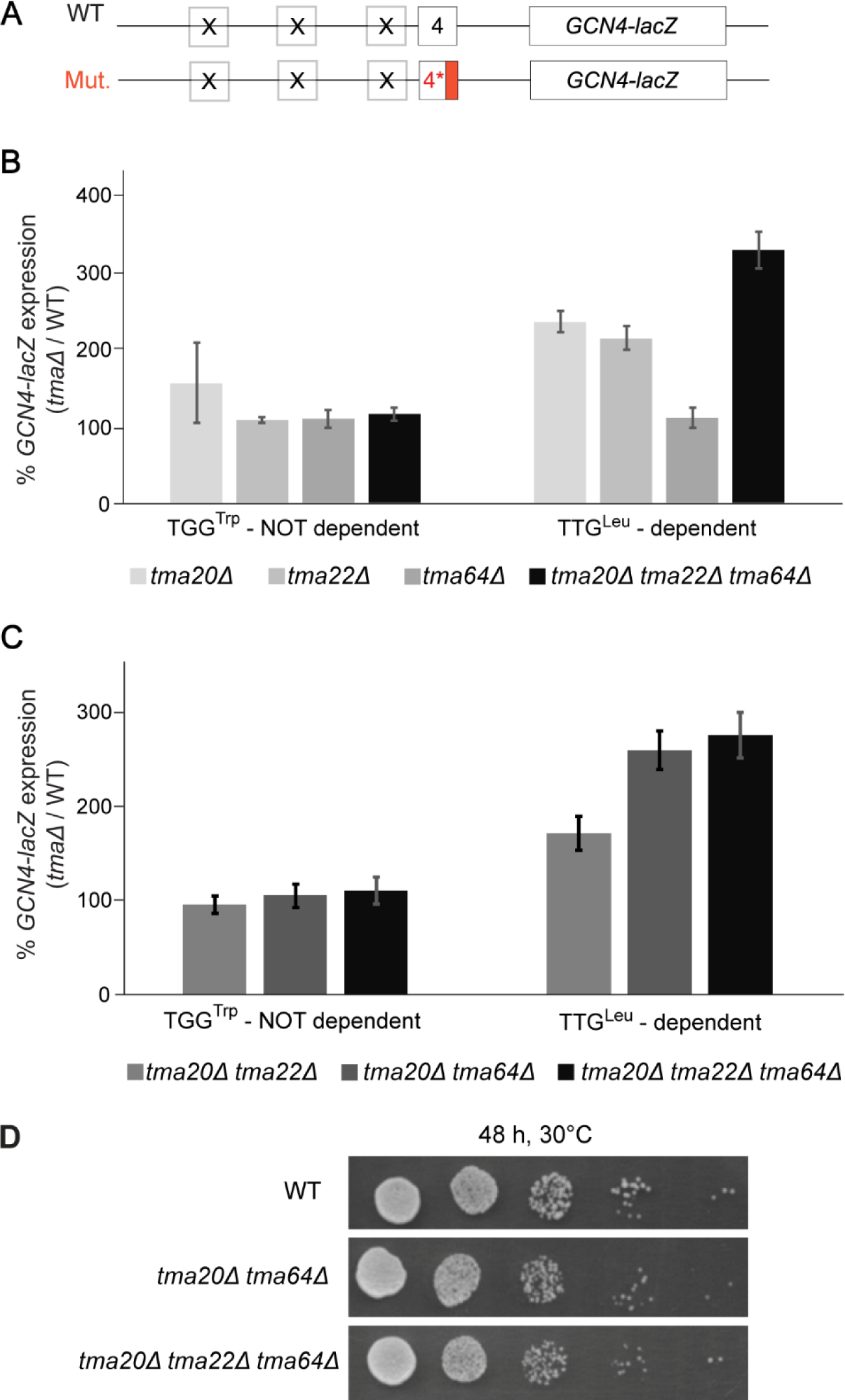
Differential effects of various *TMA* deletions on expression of uORF4-only *GCN4-lacZ* reporters equipped with Tma-independent or-dependent penultimate codons. **A)** Schematic of the uORF4-only *GCN4-lacZ* reporters harboring either WT or mutant penultimate codons. **B)** Yeast strains YSG181 (*tma20Δ*), YSG184 (*tma22Δ*), YSG178 (*tma64Δ*), or YKJ3 (*tma20Δtma22Δtma64Δ*), deleted for one or all three *TMA* genes and the corresponding WT strain YSG142 (WT BY4741) were transformed with uORF4-only reporters containing WT uORF4 (p226) or uORF4 variants with the penultimate codon CCG exchanged for Tma-dependent TTG^Leu^ (pKJ34) or Tma-independent TGG^Trp^ triplet (pKJ36). Reporter activity was assayed in whole cell extracts for at least 4 independent transformants and activities in the *tmaΔ* transformants were normalized as described in METHODS. Data are represented as ratios of normalized mean values ± SD of β-galactosidase activities in *tmaΔ* strains/WT strain. Each experiment was repeated at least once and a representative experiment is presented here. **C)** Same analysis as in (B) except comparing transformants of the double mutants YKJ6 (*tma20Δtma22Δ*) and YSG196 (*tma20Δtma64Δ*) to the triple mutant YKJ3 (*tma20Δtma22Δtma64Δ*). **D)** Serial dilutions of strains of the indicated genotypes from (C) were spotted on minimal SD medium and grown for 48 h at 30°C.

The results on the TTG^Leu^ reporter in Fig. 3B further suggest that Tma64 cannot fully substitute for the Tma20/Tma22 heterodimer, but that Tma64 does still contribute to inhibiting REI in cells lacking the heterodimer, as eliminating all three Tma proteins in the triple mutant conferred significantly greater derepression of the TTG^Leu^ reporter compared to the *tma20Δ* or *tma22Δ* single mutants (Fig. 3B). This inference was confirmed by our finding that the triple mutant and *tma20Δtma64Δ* double mutant, in which both the heterodimer and Tma64 are missing, exhibit greater derepression of the TTG^Leu^ reporter compared to the *tma20Δtma22Δ* double mutant still containing Tma64 (Fig. 3C). Consistent with this, no discernible difference in growth rate was observed between the *tma20Δtma64Δ* double mutant and the triple mutant, both of which grow slightly more slowly than the WT (Fig. 3D). Accordingly, in subsequent experiments, we considered the *tma20Δtma64Δ* and *tma20Δtma22Δtma64Δ* strains to be equivalent in completely lacking the functions of both the heterodimer and Tma64 in regulating REI.

### Tma proteins suppress REI at uORF4 variants containing Tma-independent as well as Tma-dependent penultimate codons

To extend our analysis of the impact of different penultimate codons on Tma-dependent 40S recycling and REI, we examined additional uORF4-only reporters carrying the three classes of penultimate codons listed in Table 1 corresponding to Tma-dependent, Tma-independent, and Tma–neutral for 40S recycling in the translatome^16^. As shown in Fig. 4A-B, the triple deletion mutation derepressed *GCN4-lacZ* expression from the reporter containing the Tma-dependent ATT^Ile^ codon similar to that observed for the Tma-dependent TTG^Leu^ codon analyzed above (∼2-2.5-fold derepression, cols. 7-8). Little to no derepression occurred for the Tma-independent TAC^Tyr^ reporter, as observed above for the Tma-independent TGG^Trp^ reporter (Fig. 4B, cols. 5-6). However, two of three Tma-neutral reporters, GCG^Ala^ and CTG^Leu^, display ∼1.8-2.5-fold derepression comparable to that given by the Tma-dependent reporters (cols. 3-4 vs. 7-8), whereas the neutral CCG^Pro^ codon of native uORF4 showed only slight (∼1.3-fold), but significant derepression (col. 2). The marked derepression seen for the Tma-neutral GCG^Ala^ and CTG^Leu^ reporters blurred the distinction between the effects of *tma* mutations on the uORF4 variants with Tma-dependent vs.-independent penultimate codons in Fig. 4B.

**FIGURE 4:**
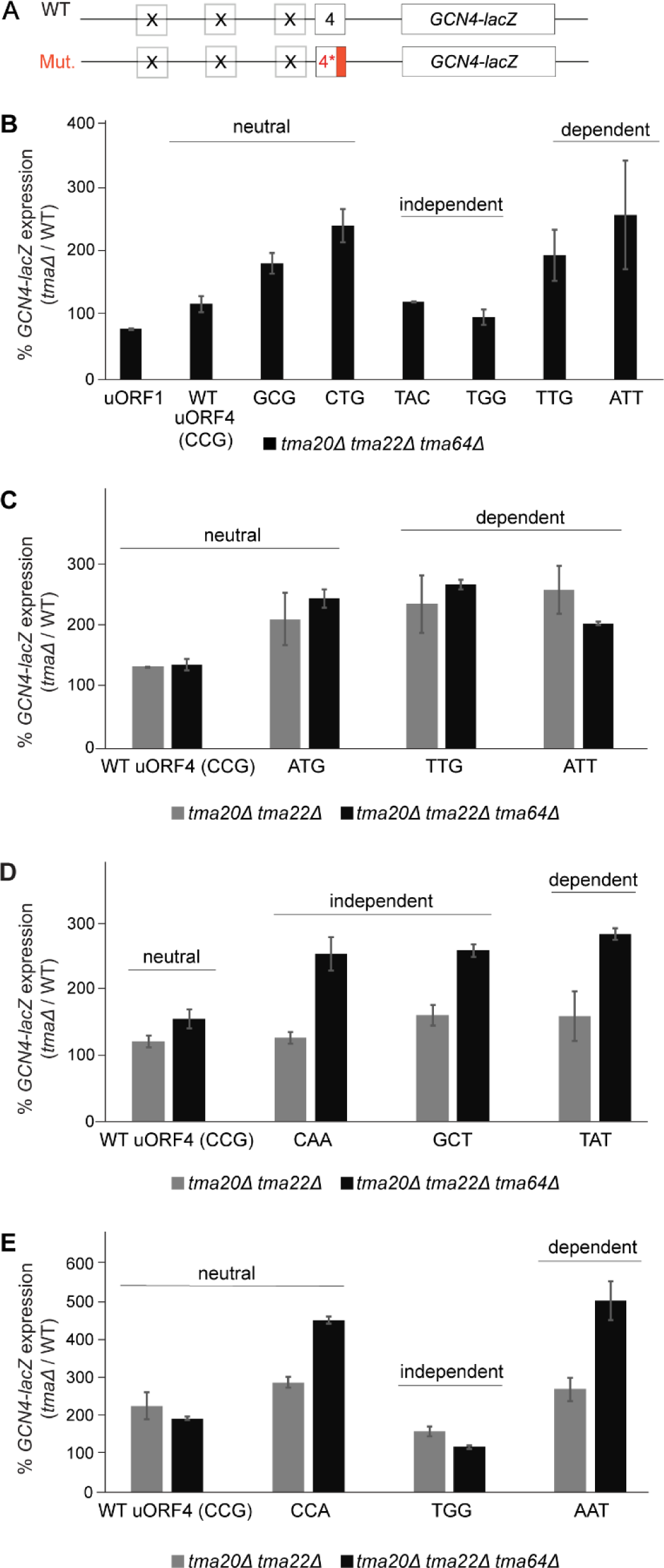
Differential effects of various *TMA* double and triple deletions on expression of uORF4-only *GCN4-lacZ* reporters equipped with Tma-independent, -neutral or -dependent penultimate codons. **A-D)** Yeast strains YKJ6 (*tma20Δtma22Δ*) and YKJ3 (*tma20Δtma22Δtma64Δ*) and the corresponding WT strain were transformed with the WT uORF4-only *GCN4-lacZ* reporter construct (p226) and variants with the uORF4 penultimate codon exchanged for the indicated neutral or Tma-dependent or -independent codons. Reporter activity was assayed in whole cell extracts for at least four independent transformants and activities in the *tmaΔ* transformants were normalized as described in METHODS. Data are represented as ratio of normalized mean values ± SD of β-galactosidase activities in *tmaΔ* strains/WT strain. Each experiment was repeated at least once, and the figure depicts representative data from one of the experiments.

**TABLE 1.** Fold-changes in uORF4-only reporter expression in *tma* mutant vs. WT strains^a^.

| Penultimate codon | *tma20*Δ*tma22*Δ /WT ratio | *tma20*Δ*tma22*Δ*tma64*Δ /WT ratio |
| --- | --- | --- |
| **Tma-neutral** |  |  |
| CCG<sup>Pro</sup> (uORF4 WT) | 1,60 | 1,78 |
| GCG<sup>Ala</sup> | 2,07 | 1,83 |
| CTG<sup>Leu</sup> | 2,35 | 2,42 |
| ATG<sup>Met</sup> | 2,64 | 2,84 |
| CCA<sup>Pro</sup> | 2,58 | 3,54 |
| **Tma-independent** |  |  |
| TACTyr | 2,11 | 1,22 |
| TGG<sup>Trp</sup> | 1,33 | 1,09 |
| CAAGln | 1,41 | 2,39 |
| GCT<sup>Ala</sup> | 2,14 | 2,73 |
| **Tma-dependent** |  |  |
| TTG<sup>Leu</sup> | 3,14 | 3 |
| ATT<sup>Ile</sup> | 3,44 | 2,85 |
| TAT<sup>Tyr</sup> | 1,52 | 2,89 |
| AAT<sup>Asn</sup> | 2,11 | 4,73 |

<sup>a</sup> Mutant uORF4-only reporters were classified according to the Tma-dependency of their penultimate codons according to Young et al. (2021). Averages of fold-changes in reporter expression from independent experiments for either the *tma20*Δ *tma22*Δ strain (YKJ6) or *tma20*Δ *tma22*Δ *tma64*Δ strain (YKJ3) versus the corresponding WT strain are tabulated.

To explore this distinction further, we examined reporters with additional penultimate codons and measured their expression in both the *tma20Δtma22Δ* double mutant and the triple mutant to provide insights into the codon dependence of Tma64 versus the Tma20/Tma22 heterodimer in suppressing REI. Considering first the results obtained in the triple mutant (Fig. 4B), an important result was that reporters with Tma-independent CAA^Gln^ or GCT^Ala^ codons (Fig. 4D, black bars) or Tma-neutral ATG^Met^ (Fig. 4C, black bars) or CCA^Pro^ codons (Fig. 4E, black bars) showed marked derepression comparable to that seen for the reporters with Tma-dependent TTG^Leu^ and ATT^Ile^ codons (Fig. 4C; black bars), TAT^Tyr^ (Fig. 4D; black bars), and AAT^Asn^ (Fig. 4E; black bars). A summary of these results in Table 1, where data from all independent experiments with the triple mutant were averaged (col. 3, *tma20Δtma22Δ tma64Δ/*WT ratio), indicates that strong derepression in the triple mutant can be observed for multiple reporters representing Tma-independent and Tma-neutral codons in addition to those containing Tma-dependent codons, with only two reporters containing Tma-independent codons TAC^Tyr^ or TGG^Trp^ being largely unaffected by elimination of all three Tma proteins.

Despite this apparent discrepancy with the previously published data on the Tma-dependency of different penultimate codons^16^, plotting the expression changes for Tma-dependent, Tma-independent or Tma-neutral penultimate codons measured in the triple mutant vs. WT (Fig. 5A) revealed a significant tendency for the four Tma-dependent codons to exhibit greater derepression in the triple mutant compared to the four Tma-independent codons tested (p=0.049). Nonetheless, since the three Tma-neutral reporters containing ATG^Met^, CTG^Leu^, or CCA^Pro^, as well as the Tma-independent GCT^Ala^ reporter did not exhibit statistically smaller derepression ratios in the triple mutant compared to the four Tma-dependent codons (Table 1, col. 3), we conclude that the Tma-dependence of penultimate codons determined previously^16^ is not a strong indicator of Tma-mediated suppression of REI following uORF4 translation.

**FIGURE 5:**
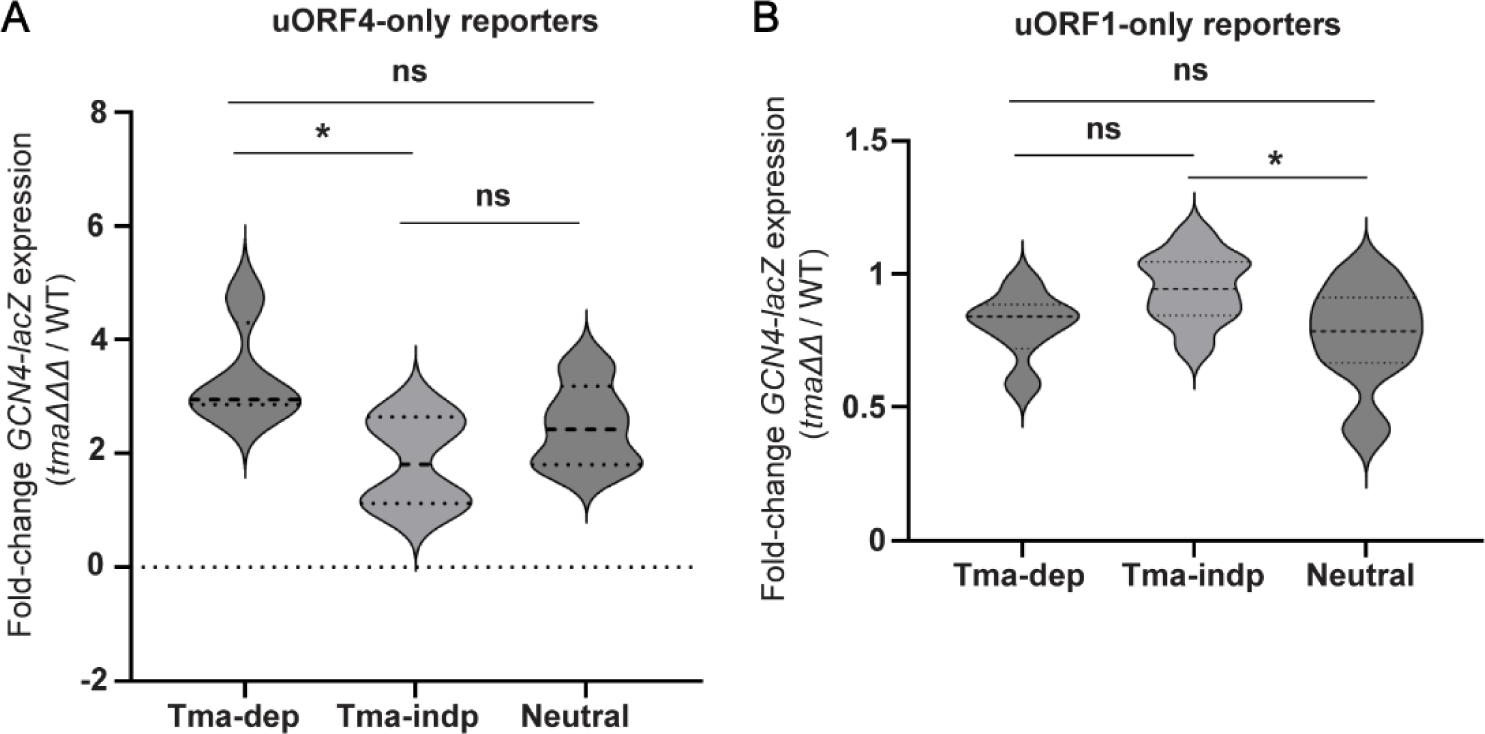
Changes in reporter expression for groups of Tma-dependent, Tma-independent or Tma-neutral penultimate codons in *tma* mutant versus WT cells. **A)** Violin plot of the average fold-changes in expression of uORF4-only *GCN4-lacZ* reporters harboring the four Tma-dependent (Tma-dep), four Tma-independent (Tma-indp) or five Tma-neutral penultimate codons in the triple deletant yeast strain YKJ3 (*tma20Δtma22Δtma64Δ*) versus isogenic WT strain, generated using the values listed in Table1 for this strain comparison. Statistical analysis was performed using two-tailed unpaired *t*-test * *p*<0.05. **B**) Violin plot of fold changes in expression of uORF1-only *GCN4-lacZ* reporters harboring the six Tma-dependent, nine Tma-independent and 14 Tma-neutral penultimate codons in *tma20Δtma64Δ* strain H4520 versus WT strain BY4741. The average fold-change of expression determined from biological replicates was calculated for each construct from the results plotted in Fig. 6. Statistical analysis was performed using the two-tailed unpaired *t*-test. * *p*<0.05.

Considering next results obtained in the *tma20Δtma22Δ* double mutant, where Tma64 can potentially substitute for the heterodimer, we found that five of the uORF4-only reporters that showed marked derepression in the triple mutant displayed substantially less derepression in the *tma20Δtma22Δ* double mutant, including reporters with CAA^Gln^, GCT^Ala^, TAT^Tyr^ (Fig. 4D), CCA^Pro^, and AAT^Asn^ penultimate codons (Fig. 4E; summarized in Table 1, cf. cols. 2-3). For these five reporters, containing Tma-independent, -neutral, or -dependent codons, it appears that Tma64 can only partially substitute for heterodimer function in suppressing REI. The Tma-independent CAA^Gln^ and Tma-dependent TAT^Tyr^ codons illustrate the most complete functional substitution by Tma64, with only slight derepression in the *tma20Δtma22Δ* double mutant despite marked derepression in the triple mutant (Fig. 4D, cols. 3-4 and 7-8, Table 1). The three reporters with ATG^Met^ (Tma-neutral) or either TTG^Leu^ or ATT^Ile^ (Tma-dependent) codons appear to represent the other extreme wherein Tma64 cannot functionally compensate and eliminating the heterodimer in the double mutant confers essentially the same degree of derepression found in the triple mutant (Fig. 4C cols. 3-8; Table 1). However, because the TTG^Leu^ reporter behaved differently in Fig. 3C in showing significantly greater derepression in the triple mutant vs. the *tma20Δtma22Δ* double mutant, it is unclear whether it belongs to this last set of reporters where Tma64 provides no functional compensation for the heterodimer. The low-level derepression observed in the triple mutant for the native uORF4 CCG^Pro^ reporter and TGG^Trp^ reporters (Fig. 4E, black bars) makes it difficult determine whether Tma64 can partially substitute for the heterodimer at these uORFs. Nevertheless, these findings overall raise the interesting possibility that the ability of Tma64 to partially substitute for the heterodimer in suppressing REI depends on the penultimate codon at uORF4, and this codon variability applies to both Tma-dependent and Tma-neutral codons. Thus, the ability of Tma64 to suppress REI in cells lacking the heterodimer does not appear to be governed by the codon-dependence of Tma factors for bulk 40S recycling^16^.

### Very few penultimate codons confer Tma-dependence at REI-permissive *GCN4* uORF1

The mammalian DENR/MCTS1 heterodimer was shown to promote REI after certain short uORFs in mammalian cells, including *ATF4* uORF1^13^; whereas the yeast Tma20/Tma22 counterpart inhibited REI following uORF translation in yeast extracts^16^. As mentioned above, it was suggested that the discrepancy between mammals and yeast arises from the inability of post-termination 40S complexes at typical yeast uORFs to remain associated with the mRNA and migrate to downstream start codons following release of the penultimate tRNA^13^ (Fig. 1B(i)(a)). To test this hypothesis, we analyzed the effects of the *tma* mutations on REI at *GCN4* uORF1. This atypical uORF is surrounded by *cis*-acting Reinitiation Promoting Elements (RPEs), some of which interact with eIF3 subunits and promote retention of post-termination 40S complexes on the mRNA, enabling high-level REI (Fig. 2). Indeed, REI after uORF1 translation is ∼20-25-fold higher than after uORF4 translation^25,26,32,33^. The retention of post-termination 40S subunits at uORF1 makes the uORF1-only *GCN4-lacZ* reporter (Fig. 6A) ideally suited to examine whether Tma factors are required to dissociate deacylated tRNA and enable reinitiation at the *GCN4* start codon in a manner limited to penultimate codons requiring the Tma factors for bulk 40S recycling in yeast^16^—in the manner observed for short uORFs in mammalian cells. If so, then uORF1-only reporters equipped with Tma-dependent codons should exhibit reduced expression in *tmaΔΔ* vs. WT cells (Fig. 1A(i)-(a)).

**FIGURE 6:**
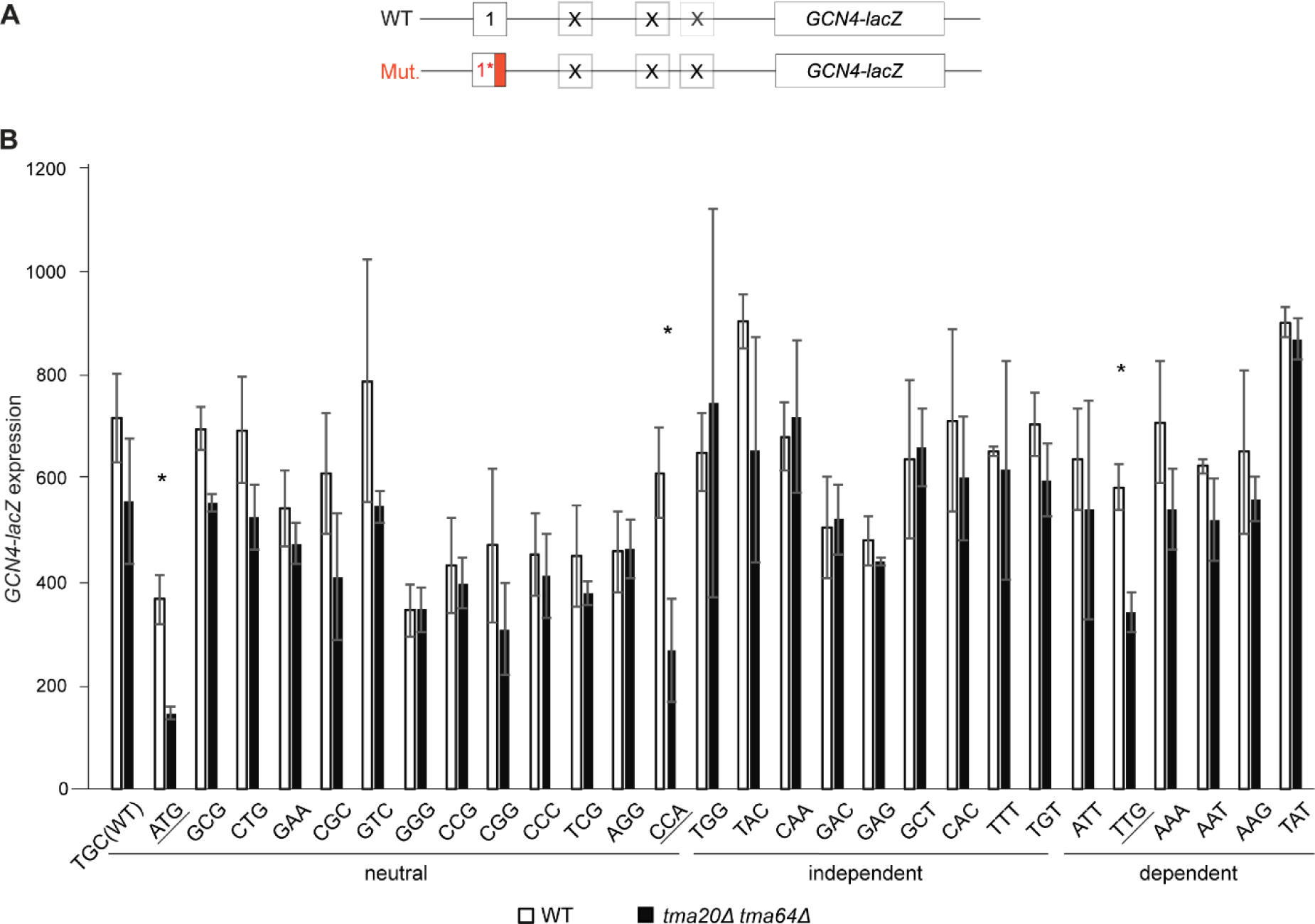
Differential effects of deleting both *TMA20* and *TMA64* on expression of uORF1-only *GCN4-lacZ* reporters equipped with neutral, Tma-independent or Tma-dependent penultimate codons. **A)** Schematic of the uORF1-only *GCN4-lacZ* reporters harboring either WT or mutant penultimate codons. **B)** Yeast strains H4520 (*tma20Δtma64Δ*) and WT strain BY4741 were transformed with uORF1-only *GCN4-lacZ* reporter constructs with uORF1 penultimate codons exchanged for the indicated neutral, Tma-independent or Tma-dependent codons. β-galactosidase activities determined for each of three independent transformants of the *tma20Δtma64Δ* strain were divided by a factor of 1.316 prior to calculating the mean activities and S.E.M. values plotted here to normalize for differences in reporter expression between the two strains that are independent of the uORF1 variants and observed for an uORF-less reporter (as described in METHODS). The mean values between mutant and WT were compared for each reporter in a two-tailed, unpaired student t-test to assign p-values (*, *p*<0.05).

We exchanged the native uORF1 penultimate codon TGC^Cys^ in the uORF1-only reporter with other penultimate codons of all three classes defined above, including six Tma-dependent, nine Tma-independent, and 13 Tma-neutral codons. The native TGC^Cys^ codon is Tma-neutral, and the native TGT^Cys^ codon of the REI-permissive uORF2 of *GCN4*^32^ was chosen as one of the Tma-independent codons. Expression of the resulting reporters was assayed in the *tma20Δtma64Δ* double mutant and WT strains (Fig. 6). At odds with the prediction that REI would be reduced in the mutant for reporters with Tma-dependent codons, only one of the six reporters of this class, containing the TTG^Leu^ codon, showed significantly reduced expression in the mutant (Fig. 6B, dependent constructs). While TTG^Leu^ was judged to be highly Tma-dependent for bulk 40S recycling, ATT^Ile^ and TAT^Tyr^ were equally so^16^, yet the reporters for the latter two codons showed unchanged expression in the mutant strain. Moreover, the constructs containing the Tma-neutral ATG^Met^ or CCA^Pro^ codons were the only reporters besides the TTG^Leu^ construct showing significantly reduced expression in *tma20Δtma64Δ* cells (Fig. 6B, neutral constructs). Plotting the expression changes for the entire set of 29 codons examined at uORF1 (Fig. 5B) revealed no significant difference in expression ratios between the Tma-dependent reporters and either the Tma-independent or Tma-neutral reporters.

Fig. 5B does reveal a significant difference between the median expression ratios between the Tma-independent and Tma-neutral groups (P = 0.0394), which reflects the slightly increased REI observed in *tmaΔΔ* cells for the Tma-independent group. None of the uORF1-only reporters, however, showed the ≥2-fold increased REI conferred by the *tma* mutations for most of the uORF4-only reporters (Fig. 5A), even for the ten cases where uORF1-only and uORF4-only reporters share the same penultimate codons (ATT, TTG, AAT, TAT, ATG, GCG, CTG, CAA, GCT, CCA). This is consistent with the prediction that the specialized RPEs at uORF1 allow 40S post-termination complexes to resist ribosome dissociation by the Tma factors that suppress REI at uORF4 lacking RPEs (Fig. 2). In summary, our findings indicate that uORF1 differs from mammalian short uORFs, including the REI-permissive uORF1 of *ATF4,* in lacking a requirement for Tma factors in dissociating deacylated tRNA from post-termination complexes at penultimate codons that impose a Tma-requirement in bulk 40S recycling. A lack of Tma-dependence on REI also applies to eight of the nine DENR-dependent codons represented among our uORF1-only reporters (ATT, GCG, CTG, CCG, CGG, CCC, AGG, and GCT). In fact, ATG is the sole DENR-dependent codon we tested that confers a dependence on Tma factors for REI at uORF1, but as noted above, ATG is Tma-neutral for 40S recycling in yeast.

In summary, neither Tma-dependence nor DENR-dependence of the penultimate codon is an accurate predictor of Tma-dependence for REI at *GCN4* uORF1 variants, despite the functional similarity between *GCN4* uORF1 and *ATF4* uORF1.

### Function of yeast ‘start-stop’ elements is not Tma-dependent

As mentioned earlier, mammalian DENR was shown to be required for REI after start-stop uORFs consisting of ATG followed by a stop codon^13,18^. To test the Tma-dependence of start-stops for downstream REI in yeast, we deleted the 2^nd^ and 3^rd^ codons from native uORF1-, uORF3-and uORF4-only constructs, without altering any other sequences, producing uStart-stop-only constructs uSt-st1, uSt-st3 and uSt-st4, respectively. (See bottom of Fig. 2 for sequences of the three uSt-st uORFs.)

First, we noticed that all three start-stop elements are more inhibitory to REI than their native uORF counterparts, reducing *GCN4-lacZ* expression in WT cells to ∼2%, ∼20%, and ∼37% of the levels observed for native uORF1-only, uORF3-only, and uORF4-only reporters, respectively (Fig. 7A). Based on findings in mammalian cells, we reasoned that converting native uORF1 to uSt-st1 would confer a dependence on REI by the Tma proteins not observed for native uORF1 with its penultimate codon TGC^Cys^ (Fig. 6, TGC(WT)), reducing expression in *tmaΔΔ* cells. This was not observed however (Fig. 7B, cols. 1-4), even though we found above that ATG^Met^ as the penultimate codon for 3-codon uORF1 conferred Tma-dependence for efficient REI (Fig. 6, ATG). Interestingly, converting uORF4 to uSt-st4 eliminated the moderate derepression observed in *tmaΔΔ* cells for the native uORF4-only reporter (Fig. 7B, cols. 9-12). This suggests that conversion to a start-stop element eliminates the requirement for Tma factors for highly efficient 40S recycling at uORF4 and repression of REI downstream. The Tma proteins appear to make no contribution to recycling and REI at native uORF3, as the uORF3-only reporter expression is unaffected by the *tma20Δtma64Δ* mutations; whereas its conversion to uSt-st3 confers a slight Tma-dependence for recycling and suppression of REI (Fig. 7, cols. 5-8). Overall, our findings suggest that, unlike their mammalian counterparts, the Tma proteins play little role in controlling REI after start-stop elements in yeast.

**FIGURE 7:**
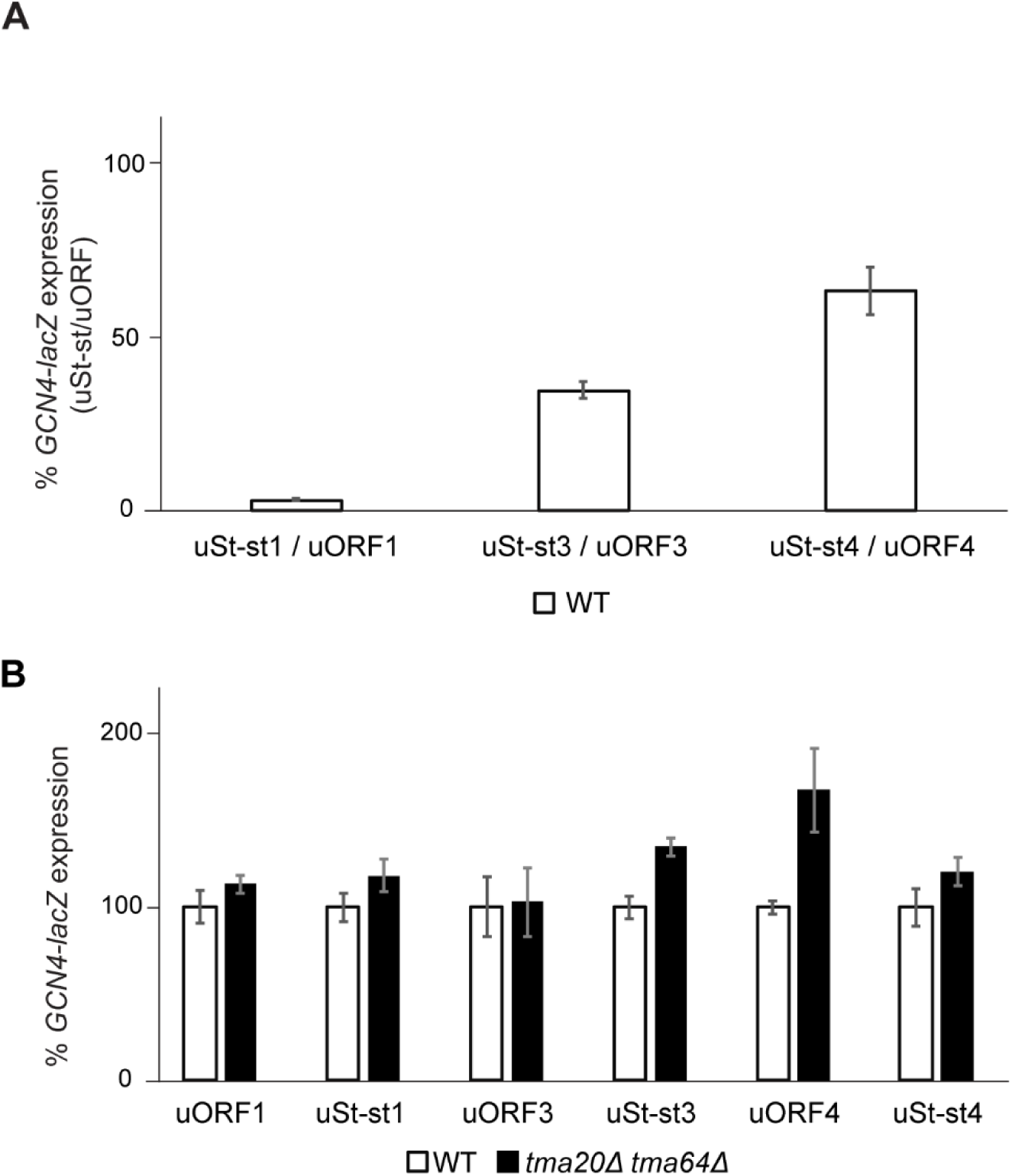
Effects of start-stop elements on downstream reinitiation in the *GCN4-lacZ* reporter system and differential effects of *TMA20* and *TMA64* deletion on re-initiation downstream of start-stop elements. **A)** WT strain BY4741 was transformed with *GCN4-lacZ* reporter plasmids containing the following single WT uORFs: uORF1-only (p209), uORF3-only (pSG61_(2)), or uORF4-only (p226); or containing single Start-stop elements uSt-st1 (pKP78), uSt-st3 (pKP76), or uSt-st4 (pKP77). β-galactosidase activities were assayed in at least 4 biological replicates and each experiment was repeated at least once, with a representative experiment depicted. Data are presented as ratios of mean reporter expression for the uSt-st reporter relative to the corresponding WT uORF reporter. **B)** Strains YSG196 (*tma20Δ tma64Δ)* and the corresponding WT were transformed with the same set of *GCN4-lacZ* reporter plasmids described in (A) and reporter expression was measured as described there. Mean reporter expression values in the *tmaΔΔ* strain were normalized as described in METHODS.

## DISCUSSION

As yeast Tma20/Tma22 and, to a lesser degree, Tma64 have been implicated in 40S ribosome recycling *in vivo*^15,16^, we used an established reporter system based on *GCN4* translational control to examine the influence of yeast Tma20/MCTS1, Tma22/DENR and Tma64/eIF2D on REI downstream of short uORFs optimized for either REI (uORF1) or for recycling and suppression of REI (uORF4)^5^, and to dissect the functional interplay between the Tma factors in regulating these alternative post-termination outcomes. We have examined the effects of eliminating the Tma factors on variants of uORF1 or uORF4 containing various penultimate codons that conferred strong dependence on Tma proteins for recycling at most stop codons in yeast, or equipped with different codons showing little dependence on Tma factors in bulk recycling.

Using the uORF4-only *GCN4-lacZ* reporter, we first demonstrated that inserting Tma-dependent codon TTG^Leu^ led to substantially increased REI in mutants lacking either *TMA20* or *TMA22* singly, but not *TMA64.* Deletion of *TMA64* in the *tma20Δtma22Δ* mutant enhanced this defect, suggesting that Tma64 can partially compensate for the absence of Tma20/Tma22 in suppressing REI. This conclusion agrees with 40S profiling data from^16^, wherein a single deletion of *TMA20* or *TMA22,* but not *TMA64*, led to accumulation of unrecycled 40S subunits at stop codons translatome-wide, and this phenotype was more pronounced in the *tma20Δtma64Δ* deletion strain lacking both the heterodimer and Tma64. Consistent with this, we observed essentially the same increased REI for the TTG^Leu^ uORF4 variant in the *tma20Δtma64Δ* double mutant and *tma20Δtma22Δtma64Δ* triple mutant. We concluded that intact Tma20/Tma22 heterodimer is indispensable for WT 40S recycling and suppression of REI but receives functional support from Tma64 at the stop codon of this uORF4 variant.

Examining a total of 12 different uORF4 variants with different penultimate codons and native uORF4 revealed that the absence of all three Tma proteins increased REI by ≥2-fold for all but three uORFs: native uORF4 containing CCG^Pro^ and the TAC^Try^ and TGG^Trp^ variants. Although none of these codons were found to confer Tma-dependence on bulk recycling^16^, six other uORF4 variants containing Tma-independent or Tma-neutral penultimate codons did increase REI by ≥2-fold in the triple mutant (CAA^Gln^, GCT^Ala^, GCG^Ala^, CTG^Leu^, ATG^Met^, and CCA^Pro^), as did all four reporters containing Tma-dependent codons (Table 1). Even though the median *tma20Δtma22Δtma64Δ*/WT derepression ratio is significantly larger for the reporters with Tma-dependent vs. Tma-independent codons (Fig. 5A), the Tma-dependence in bulk 40S recycling determined previously^16^ is generally a weak predictor of Tma-dependence in promoting 40S recycling and suppressing REI for different penultimate codons at uORF4 (Table 1). Recycling and REI at the uORF4 variant containing the Tma-independent TGG^Trp^ codon appeared to be the least dependent on Tma factors among all 13 different penultimate codons we examined. While the underlying reason remains unknown, it is interesting that TGG^Trp^ is the sole codon for tryptophan.

Our finding that the Tma proteins frequently inhibit REI at uORF4 variants containing Tma-neutral or Tma-independent codons is difficult to explain if dissociation of the empty 40S subunit occurs spontaneously or is catalyzed by other factors at these penultimate codons, which would suppress REI in the presence or absence of Tma factors (Fig. 1B(ii) (a)). Hence, in the model proposed in Fig. 1B(ii) (b), we suggest that the Tma factors do function in dissociating the empty 40S subunit at Tma-independent codons (recycling step 2 in WT), such that retention of the 40S subunit in *tmaΔΔ* cells allows REI downstream by the fraction of 40S subunits where release of the deacylated tRNA occurred independently of Tma factors to confer increased REI (Fig. 1B(ii) (b), *tmaΔΔ*). Our observation that the Tma proteins also inhibit REI at uORF4 variants containing Tma-dependent codons is likewise difficult to explain because the failure to release the deacylated tRNA in the *tmaΔΔ* mutant should block REI even though Tma-catalyzed dissociation of the 40S subunit from mRNA is absent, thus keeping REI low in *tmaΔΔ* cells (Fig. 1B(i) (a)). Therefore, it seems necessary to propose that the deacylated tRNA eventually dissociates spontaneously in the absence of Tma factors, so that loss of Tma-catalyzed dissociation of the empty 40S subunit then enables REI, effecting the increased REI observed in *tmaΔΔ* cells (Fig. 1B(i) (b), just as proposed above for Tma-independent penultimate codons (Fig. 1B(ii) (b)).

The model proposed in Fig. 1B(ii)-(b) may seem problematic in positing that 40S subunit dissociation is catalyzed by the Tma factors at codons judged to be Tma-independent for bulk 40S recycling. However, perhaps Tma-independent recycling can operate at all stop codons, at least in *tmaΔΔ* mutants, and recycling at Tma-dependent penultimate codons simply relies more heavily on Tma factors than at Tma-independent codons. This possibility is consistent with the fact that eliminating all three Tma proteins confers only a small effect on cell growth, implying the frequent occurrence of Tma-independent recycling throughout the translatome. Another possibility is that recycling at Tma-dependent penultimate codons relies heavily on the Tma factors for both steps of recycling, whereas the first step (deacylated tRNA release) occurs spontaneously, or is catalyzed by other recycling factors, at Tma-independent codons.

Comparing the derepressing effects of the *tma20Δtma22Δtma64Δ* triple mutation to the *tma20Δtma22Δ* double mutation for the set of uORF4-only reporters allowed us to assess the penultimate codon-dependence for recycling carried out by Tma64 alone. Interestingly, we found evidence that the contribution of Tma64 to recycling and suppressing REI appears to vary for different penultimate codons.

We also investigated whether the Tma factors function similarly to MCTS1/DENR in promoting, rather than impeding, REI at short uORFs that are permissive for REI owing to retention of eIFs during uORF translation. It was proposed that the persistence of eIFs on 40S post-termination complexes prevents dissociation of the 40S subunits following MCTS1/DENR-catalyzed release of deacylated tRNA, allowing resumption of 40S migration and rebinding of TC for reinitiation downstream. This explained why REI was impaired on depletion of MCTS1/DENR only for uORFs containing DENR-dependent penultimate codons^13^ (Fig. 1A(i)-(a) and (ii)-(a)). To investigate this mechanism in yeast, we examined the *GCN4* uORF1-only reporter in which RPEs surrounding uORF1 increase eIF3 and eIF4F occupancy of the post-termination 40S subunits and confer high-frequency REI at *GCN4* (Fig. 2)^25,26,32,33^. Consistent with results for MCTS1/DENR, REI was not reduced in the *tmaΔΔ* mutant for native uORF1 nor for 21 other uORF1 variants equipped with Tma-neutral or Tma-independent penultimate codons, but was reduced when uORF1 contained the Tma-dependent TTG^Leu^ codon. However, REI was not reduced in *tmaΔΔ* cells at five other uORF1 variants with Tma-dependent codons. This included the TAT codon present in uORF1 of *ATF4* mRNA—functionally equivalent to *GCN4* uORF1^38^—that confers DENR-dependent REI on *ATF4* mRNA^13^. Moreover, REI was markedly reduced in *tmaΔΔ* cells when uORF1 contained the Tma-neutral penultimate codons ATG and CCA. Thus, dependence on the Tma factors for REI was conferred on uORF1 by only three of 29 penultimate codons examined, only one of which (TTG) is Tma-dependent for bulk 40S recycling^16^. Moreover, there was no significant difference in median *tmaΔΔ* derepression ratios between uORF1-only reporters containing Tma-dependent vs. Tma-neutral or Tma-independent codons (Fig. 5B). Overall, it appears that the Tma factors do not promote REI at uORF1 in the manner described for MCTS1/DENR at REI-permissive uORFs in mammalian cells.

One possibility is that the specialized interaction of *GCN4* uORF1 RPEs with eIF3 generally overrides the requirement for Tma factors in dissociating the deacylated tRNA at Tma-dependent penultimate codons. By preventing 40S dissociation from the mRNA, eIF3 might provide sufficient time for dissociation of the deacylated tRNA, spontaneously or by a Tma-independent pathway, to prevent reduced REI in *tmaΔΔ* cells, as described in the model of Fig. 1A(i)-(b). In support, numerous tRNAs were found to readily dissociate from the 40S subunit without the help of any dedicated factor ^19^. Furthermore, structural analysis of MCTS1/DENR bound to the 40S subunit indicates that MCTS1 might occupy the same binding site on the ribosome as the eIF3a, b, and c subunits of mammalian eIF3^39^. In addition, the ribosome binding interface in MCTS1 seems to be well conserved from yeast to higher eukaryotes (red dots in Fig. 8). This might indicate competitive ribosome binding between eIF3 and Tma20/Tma22, such that the uORF1 RPEs would promote binding of eIF3a vs. the Tma heterodimer and thereby prevent the latter from influencing REI at uORF1.

**FIGURE 8:**
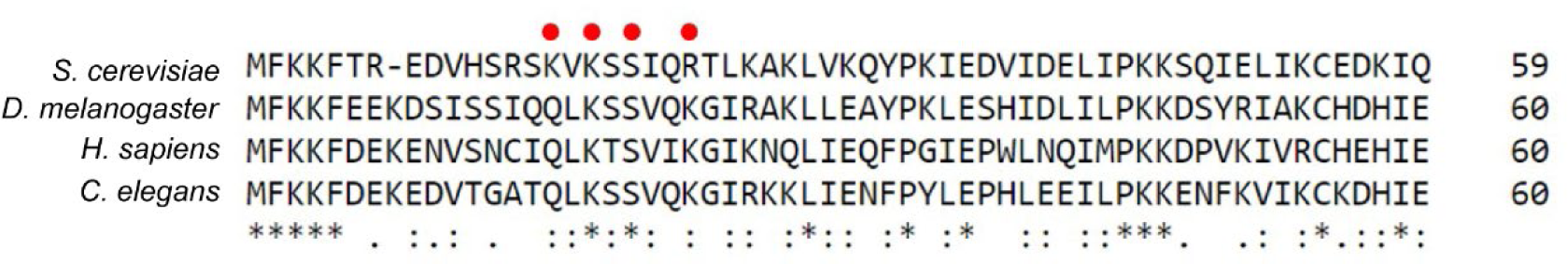
Alignment of *S. cerevisiae* (yeast), *D. melanogaster* (fly), *H. sapiens* (human) and *C. elegans* (worm) Tma20/MCTS1 proteins. Multiple sequence alignment was performed using Clustal Omega. This alignment illustrates the conservation of amino acid residues of the ribosome binding interface described previously^39^ and marked here by red dots.

Presumably, interaction of deacylated leucyl tRNA with the TTG^Leu^ codon is too stable to overcome the requirement for Tma-mediated release, leading to the observed reduced REI for the TTG^Leu^ uORF1 variant in *tmaΔΔ* cells, in the manner predicted for DENR-dependent codons (Fig. 1A(i) (a)). The same might be true for interaction of elongator methionyl tRNA and prolyl tRNA with their cognate ATG^Met^ and CCA^Pro^ codons, which also displayed Tma-dependence for REI at uORF1, even though ATG and CCA were not identified as being Tma-dependent for bulk 40S recycling^16^. Supporting this idea, ATG was identified as a DENR-dependent penultimate codon in mammalian cells both in bulk 40S recycling and reporter REI assays^13^.

Another consequence of introducing ATG as the penultimate codon of *GCN4* uORF1 not yet mentioned is a significant reduction in REI in WT cells (P = 0.04), which is further exacerbated in the *tmaΔΔ* mutant (Fig. 6). A similar reduction in WT cells was found for the GGG^Gly^ codon (P = 0.03) but was not exacerbated in *tmaΔΔ* cells (Fig. 6B, ATG, WT vs. *tmaΔΔ*). Somewhat smaller reductions also occurred only in WT cells with the CCG^Pro^, CCC^Pro,^ AGG^Arg^ and GAG^Glu^ codon replacements (P = 0.09) (Fig. 6B). CCG^Pro^ is the evolutionarily conserved penultimate codon at uORFs 3 and 4 and both CCG^Pro^ and CCC^Pro^ were observed previously to inhibit REI when introduced at uORF1^33^. This last effect might arise from slow decoding or termination at the adjacent stop codon in the presence of these penultimate proline codons^33^.

In addition to *ATF4* uORF1 and other REI-permissive uORFs, the MCTS1/DENR heterodimer was found to promote REI after translation of the *ATF4* start-stop element preceding uORF1^13^ and at various other start-stop elements in optimum Kozak context^13,17,18,20^. In contrast, we found no evidence for Tma-dependent recycling and REI at the start-stop elements we derived from *GCN4* uORFs 1, 3, or 4, all of which contain the optimum A-rich sequence context for initiation in yeast. Converting uORF3 and uORF4 to start-stop elements decreased REI in WT cells and essentially eliminated the increased REI observed for native uORF4 in *tmaΔΔ* cells. It appears that uSt-st4 exhibits an elevated level of spontaneous 40S recycling that bypasses the contribution of Tma factors to recycling seen at native uORF4. Converting uORF1 to start-stop element uSt-st1 also dramatically decreased REI in WT cells, apparently disabling the RPEs that enable high-level REI at native uORF1. Although uSt-st1 behaves like a typical inhibitory uORF, 40S recycling appears to occur without any contribution from the Tma factors at this element, as also observed for sSt-st4. The fact that uSt-st1, uSt-st4, and uSt-st3 do not show decreased REI in *tmaΔΔ* cells in the manner we observed for the *GCN4* uORF1 variant with ATG^Met^ as penultimate codon (Fig. 6B) is not surprising considering that, similar to uORF4, the St-st elements appear to lack the capacity for eIF3-mediated retention of 40S post-termination complexes (Fig. 2).

In summary, we found that most replacements of the uORF4 penultimate codon confer increased REI in mutants lacking both Tma64 and the Tma20/Tma22 heterodimer compared to the native CCG codon, suggesting a codon-dependent role for the Tma factors in dissociating 40S post-termination complexes at this uORF, even when the penultimate triplets were shown previously to be Tma-independent for recycling at other stop codons throughout the translatome. These findings imply that the deacylated tRNA can dissociate in the absence of Tma factors at both Tma-dependent and Tma-independent penultimate codons, so that loss of Tma-catalyzed recycling of the empty 40S subunit enables elevated REI at uORF4 in the mutant cells. In contrast to findings on short, REI-permissive uORFs in mammalian cells, and *ATF4* uORF1 in particular^13^, we found that only one of six Tma-dependent penultimate codons introduced at *GCN4* uORF1 conferred reduced REI in *tma20Δtma64Δ* cells. These findings indicate that dissociation of the deacylated tRNA and subsequent reinitiation generally can proceed without Tma factors at uORF1, even at penultimate codons where these proteins enhance bulk 40S recycling. Together, our results on uORF1 and uORF4 account for our previous finding that *GCN4* translational control was essentially unperturbed in mutants lacking both Tma20/Tma22 and Tma64^29^—in stark contrast to the requirement for MCTS1/DENR for induction of *ATF4* mRNA translation. Also at odds with findings on mammalian MCTS1/DENR, converting uORF1 to a start-stop element did not confer Tma-dependent REI, and converting uORF4 to a start-stop element decreased REI and diminished the contribution of Tma factors in suppressing REI. Overall, the influence of Tma factors on REI at short regulatory uORFs in yeast cells differs considerably from that observed for mammalian MCTS1/DENR.

## MATERIALS AND METHODS

### Yeast strains and plasmids

The genotypes of all yeast strains employed in this study are listed in Table S1. Yeast mutants lacking one or more *TMA* genes were constructed using DNA cassettes bearing *kanMX4*, *hphNT1* or *natNT2* resistance markers contained in plasmids pFA6a-KanMX4^40^ (acquired from J.H. Hegeman), pZC3 and pZC4^41^, respectively. The cassettes were amplified by PCR using gene-specific primers SG325 and SG326 (*TMA64*/YDR117C) and KJ1 and KJ2 (*TMA22*/YJR014W) (see Table S3 for complete list of primers). Subsequently, the amplified DNA was used to transform yeast to confer resistance to antibiotics Geneticin G418 (BioConcept), Nurseothricine (Jena Bioscience), or Hygromycin B (Carl Roth GmbH), respectively, on YPD agar medium. YSG178 (*tma64Δ*) was derived from WT BY4741 (YSG142) by transformation with a *TMA64*-specific *kanMX4* deletion cassette. YSG196 (*tma20Δtma64Δ*) was generated from YSG181 (*tma20Δ*) by transformation with a *TMA64*-specific *natNT2* deletion cassette. YKJ3 (*tma20Δtma22Δtma64Δ*) was generated from YSG196 (*tma20Δtma64Δ*) by transformation with a *TMA22*-specific *hphNT1* cassette. YKJ6 (*tma20Δtma22Δ*) was generated from YSG181 (*tma20Δ*) by transformation with a *TMA22*-specific *natNT2* cassette. Deletion of WT alleles and replacement with the appropriate deletion cassettes was verified by colony PCR^42^ using gene-specific primers SG295 (for *TMA20*/YER007C-A), SG294 (for *TMA64*/YDR117C deletions) and SG296 (for *TMA22*/YJR014W deletions) in combination with universal primer PB238 complementary to all deletion cassettes. Strains YSG181 (*tma20Δ*) and YSG184 (*tma22Δ*) were acquired from the Euroscarf yeast deletion collection.

All plasmids employed in this study are listed in Table S2. All uORF4-only reporters were created by replacing the *SalI*-*BstEII* fragment of uORF4-only *GCN4-lacZ* construct p226^43^ with the appropriate *SalI*-*BstEII* fragments generated by either PCR primer mutagenesis or DNA synthesis. Fragments generated by PCR primer mutagenesis were amplified using common forward primer KJ27 (mapping near the *SalI* cloning site) and the following mutagenic reverse primers (mapping near the *BstEII* site) for constructing the indicated plasmids: primer KJ25 for plasmid pKJ34, primer KJ26 for pKJ35, primer KJ24 for pKJ36, primer KJ78 for pKJ57, primer KJ80 for pKJ58, primer KJ76 for pKJ59, primer KJ74 for pKJ61, primer KJ77 for PKJ62. DNA fragments for constructing pKJ17, pKJ23, pKJ20 and pKJ21 were synthesized by GeneArt Gene Synthesis (Thermo Fisher). All fragment inserts in the final constructs were verified by Sanger sequencing, and the DNA sequences are listed in Table S4.

The start-stop constructs pKP76, pKP77 and pKP78 were created by replacing the *SalI-BstEII* fragment of the *GCN4-lacZ* construct pSG194^33^; a uORF1-only construct created by replacing the coding sequence of uORF1 by that of uORF2) by the appropriate *SalI-BstEII* fragments generated by GeneArt Gene Synthesis (Thermo Fisher). These fragments contain point mutations in start codons of uORFs (point mutations in uORFs 1, 2 and 4 in pKP76, point mutations in uORFs 1, 2 and 3 in pKP77 and point mutations in uORFs 2, 3 and 4 in pKP78) and deletions of sense codons between start and stop codons in uORF3 (pKP76), uORF4 (pKP77) and uORF1 (pKP78), respectively, which creates start-stop elements uSt-st3, uSt-st4 and uSt-st1. The mutation scheme of these constructs is also shown in Fig. 2. All fragment inserts in the final constructs were verified by Sanger sequencing, and the DNA sequences are listed in Table S4.

All uORF1-only reporters were created by replacing the *SalI*-*BstEII* fragment of WT *GCN4-lacZ* construct p180^44^ with the appropriate *SalI*-*BstEII* fragments, generated by DNA synthesis by LifeSct LLC, containing point mutations in the start codons of uORFs 2-4 (*uORF2* ATG to CTG, *uORF3* ATG to AGG, and *uORF4* ATG to AGG) and either WT uORF1 (plasmid pSG61) or uORF1 variants with altered third codons (pSG62-pSG89). The inserts in the resulting plasmids were sequenced in their entirety to confirm the desired sequences. The sequence of the *SalI*-*BstEII* fragment of pSG61 (with WT uORF1) is listed in Table S4; sequences of all other constructs differ only by the replacements of the third uORF1 codon with the triplet indicated in Table S2 for the corresponding plasmids.

### *GCN4-lacZ* reporter assays

β-galactosidase specific activities (in units of nmol of ONPG cleaved per min per mg of protein) were assayed in whole cell extracts as described previously^33^ in yeast transformants of uORF4-only and uSt-st *GCN4-lacZ* reporter plasmids, and as previously described^45^ for transformants of uORF1-only reporter plasmids and the “uORF-less” *GCN4-lacZ* reporter p227^30^.

For the uORF4-only and uSt-st-only reporter assays, the normalized mean ± SD of β-galactosidase activities for each reporter construct in transformants of *tmaΔ* strains was determined by dividing the mean and SD of unnormalized β-galactosidase activities by a normalization factor. This factor was calculated separately for each individual experiment by assaying the activity of the “uORF-less” reporter (p227) in WT and *tmaΔ* strains in 4 transformants under the same conditions as uORF4-only/uSt-st-only reporters. Then, the mean of *tmaΔ* “uORF-less” activities was divided by the mean of WT “uORF-less” activities, determining the normalization factor.

For the uORF1-only reporter assays, β-galactosidase activities determined for each replicate transformant of the *tma20Δtma64Δ* strain were divided by a factor of 1.316 prior to calculating the mean activities to be compared to the mean unnormalized activities for the replicate transformants of the WT strain in a two-tailed, unpaired student t-test to assign p-values to differences between the means in mutant vs. WT cells. The normalization factor was determined by assaying 6 transformants each of the same *tma20Δtma64Δ* and WT strains harboring the “uORF-less” reporter (p227) under the same conditions employed for uORF1-only reporters. The *tma20Δtma64Δ/*WT ratio of mean expression values was 1.316, which differed significantly from unity in a two-tailed t-test with a p-value of 0.003.

### Yeast spotting assay

Yeast strains were spotted onto minimal synthetic defined (SD) medium plates in five serial 10-fold dilutions (starting with OD600 0.5) and grown for 48 h at 30°C.

### Multiple sequence alignment

Multiple sequence alignment of *S. cerevisiae* Tma20 (UniProt accession P89886), *D. melanogaster* MCTS1 (UniProt (UniProt accession Q9W445), *C. elegans* C11D2.7 (UniProt accession Q8MXH7) and *H. sapiens* MCTS1 accession Q9ULC4) protein sequences was performed using Clustal Omega^46^ with default settings through European Bioinformatics Institute Tools services^47^.

### Statistical analysis

The normality of all datasets visualized in the violin plots in Fig. 5 was tested by the Shapiro-Wilks test and then a two-tailed unpaired *t*-test was used to test differences between two experimental groups. Derived *p*-values < 0.05 were considered statistically significant. Statistical analysis and visualization were performed in GraphPad Prism, version 9.4.1 (GraphPad Software).

## ACKNOWLEDGEMENTS

We are thankful to all members of our laboratories for fruitful discussions. This work was supported in part by a Grant of Excellence in Basic Research (EXPRO 2019) provided by the Czech Science Foundation (19-25821X), the Praemium Academiae grant provided by the Czech Academy of Sciences, CZ.02.01.01/00/22_008/0004575 RNA for therapy by ERDF and MEYS (all to L.S.V.), by GA UK project no. 339022 by Charles University Grant Agency (to K.J.), and by the Intramural Program of the National Institutes of Health (S.G. and A.G.H).

## SUPPLEMENTARY DATA

Supplementary Data are available online.

## SUPPLEMENTARY TABLES

**Table S1.** Yeast strains used in this study. This table lists all yeast strains employed in this study along with their respective parental strains used to generate them. The details of strain constructions are described in MATERIALS AND METHODS.

| **Name:** | **Genotype:** | **Parental strain:** | **Source:** |
| --- | --- | --- | --- |
| YSG142 (WT BY4741) | *MATa his3Δ1 leu2Δ0 met15Δ0 ura3Δ0* | - | Euroscarf |
| YSG178 ($\Delta tma64$) | *MATa his3Δ1 leu2Δ0 met15Δ0 ura3Δ0 ydr117CΔ::kanMX4* | YSG142 | This study |
| YSG181 ($\Delta tma20$) | *MATa his3Δ1 leu2Δ0 met15Δ0 ura3Δ0 yer007C-AΔ::kanMX4* | YSG142 | Euroscarf |
| YSG184 ($\Delta tma22$) | *MATa his3Δ1 leu2Δ0 met15Δ0 ura3Δ0 yjr014WΔ::kanMX4* | YSG142 | Euroscarf |
| YSG196 ($\Delta tma20 \Delta tma64$) | *MATa his3Δ1 leu2Δ0 met15Δ0 ura3Δ0 yer007C-AΔ::kanMX4 ydr117CΔ::natNT2* | YSG181 | This study |
| YKJ3 ($\Delta tma20 \Delta tma22 \Delta tma64$) | *MATa his3Δ1 leu2Δ0 met15Δ0 ura3Δ0 yer007C-AΔ::kanMX4 ydr117CΔ::natNT2 yjr014WΔ::hphNT1* | YSG196 | This study |
| YKJ6 ($\Delta tma20 \Delta tma22$) | *MATa his3Δ1 leu2Δ0 met15Δ0 ura3Δ0 yer007C-AΔ::kanMX4 yjr014WΔ::natNT2* | YSG181 | This study |
| BY4741 | *MATa his3Δ1 leu2Δ0 met15Δ0 ura3Δ0* | - |  |
| H4520 | *MATa his3Δ1 leu2Δ0 met15Δ0 ura3Δ0 ydr117CΔ::hygMX4; yer007C-AΔ::kanMX4* | BY4741 | <sup>1</sup> |

**Table S2.** Plasmids used in this study. This table lists all plasmids used in this study, with construction details described in MATERIALS AND METHODS.

| **Plasmid:** | **Description:** | **Reference:** |
| --- | --- | --- |
| p227 (uORF-less) | low copy *URA3 GCN4-lacZ* plasmid containing the uORF-less *GCN4* leader created by point mutations in uORF1 (*HindIII*), uORF2 (*EcoRI*), uORF3 (*KpnI*) and uORF4 (*BglII*) | <sup>2</sup> |
| p226 (uORF4-only) | low copy *URA3 GCN4-lacZ* plasmid containing *GCN4* leader with uORF4 only at its original position | <sup>2</sup> |
| pKJ34 | p226 with uORF4 penultimate codon mutated to TTG | This study |
| pKJ35 | p226 with uORF4 penultimate codon mutated to ATT | This study |
| pKJ36 | p226 with uORF4 penultimate codon mutated to TGG | This study |
| pKJ57 | p226 with uORF4 penultimate codon mutated to CAA | This study |
| pKJ58 | p226 with uORF4 penultimate codon mutated to GCT | This study |
| pKJ59 | p226 with uORF4 penultimate codon mutated to TAT | This study |
| pKJ61 | p226 with uORF4 penultimate codon mutated to AAT | This study |
| pKJ62 | p226 with uORF4 penultimate codon mutated to CCA | This study |
| pKJ23 | p226 with uORF4 penultimate codon mutated to ATG | This study |
| pKJ20 | p226 with uORF4 penultimate codon mutated to TAC | This study |
| pKJ21 | p226 with uORF4 penultimate codon mutated to GTG | This study |
| pKP76 | low copy *URA3 GCN4-lacZ* plasmid containing uORF3 only at its original position mutated into a start-stop element | This study |
| pKP77 | low copy *URA3 GCN4-lacZ* plasmid containing uORF4 only at its original position mutated into a start-stop element | This study |
| pKP78 | low copy *URA3 GCN4-lacZ* plasmid containing uORF1 only at its original position mutated into a start-stop element | This study |
| pSG194 | low copy *URA3 GCN4-lacZ* plasmid containing an an uORF1 variant only with coding sequence replaced by the corresponding sequence of uORF2 | 3 |
| pKJ17 | p226 with uORF4 penultimate codon mutated to CTG | This study |
| pFA6a-KanMX4 | pFA6 backbone containing *KanMX4* resistance marker-based deletion cassette | 4 |
| pZC3 | pFA6 backbone containing *hphNT1* resistance marker-based deletion cassette | 5 |
| pZC4 | pFA6 backbone containing *natNT2* resistance marker-based deletion cassette | 5 |
| pSG61 | low copy *URA3 GCN4-lacZ* plasmid containing the *GCN4* leader with uORF1 only at its original position | This study |
| pSG62 | pSG61 with uORF1 penultimate codon mutated to ATT | This study |
| pSG63 | pSG61 with uORF1 penultimate codon mutated to TTG | This study |
| pSG64 | pSG61 with uORF1 penultimate codon mutated to AAA | This study |
| pSG65 | pSG61 with uORF1 penultimate codon mutated to AAT | This study |
| pSG66 | pSG61 with uORF1 penultimate codon mutated to AAG | This study |
| pSG67 | pSG61 with uORF1 penultimate codon mutated to TAT | This study |
| pSG68 | pSG61 with uORF1 penultimate codon mutated to ATG | This study |
| pSG69 | pSG61 with uORF1 penultimate codon mutated to GCG | This study |
| pSG70 | pSG61 with uORF1 penultimate codon mutated to CTG | This study |
| pSG71 | pSG61 with uORF1 penultimate codon mutated to TGG | This study |
| pSG72 | pSG61 with uORF1 penultimate codon mutated to TAC | This study |
| pSG73 | pSG61 with uORF1 penultimate codon mutated to CAA | This study |
| pSG74 | pSG61 with uORF1 penultimate codon mutated to GAC | This study |
| pSG75 | pSG61 with uORF1 penultimate codon mutated to GAG | This study |
| pSG76 | pSG61 with uORF1 penultimate codon mutated to TGT | This study |
| pSG77 | pSG61 with uORF1 penultimate codon mutated to GCT | This study |
| pSG78 | pSG61 with uORF1 penultimate codon mutated to CAC | This study |
| pSG79 | pSG61 with uORF1 penultimate codon mutated to TTT | This study |
| pSG80 | pSG61 with uORF1 penultimate codon mutated to GAA | This study |
| pSG81 | pSG61 with uORF1 penultimate codon mutated to CGC | This study |
| pSG82 | pSG61 with uORF1 penultimate codon mutated to GTC | This study |
| pSG83 | pSG61 with uORF1 penultimate codon mutated to GGG | This study |
| pSG84 | pSG61 with uORF1 penultimate codon mutated to CCG | This study |
| pSG85 | pSG61 with uORF1 penultimate codon mutated to CGG | This study |
| pSG86 | pSG61 with uORF1 penultimate codon mutated to CCC | This study |
| pSG87 | pSG61 with uORF1 penultimate codon mutated to TCG | This study |
| pSG88 | pSG61 with uORF1 penultimate codon mutated to AGG | This study |
| pSG89 | pSG61 with uORF1 penultimate codon mutated to CCA | This study |
| pSG61 (2)<br>(uORF3-only) | low copy *URA3 GCN4-lacZ* plasmid containing the *GCN4* leader with uORF3 only at its original position | <sup>6</sup> |
| p209 (uORF1-only) | low copy *URA3 GCN4-lacZ* plasmid containing the *GCN4* leader with uORF1 only at its original position | <sup>7</sup> |

**Table S3.**
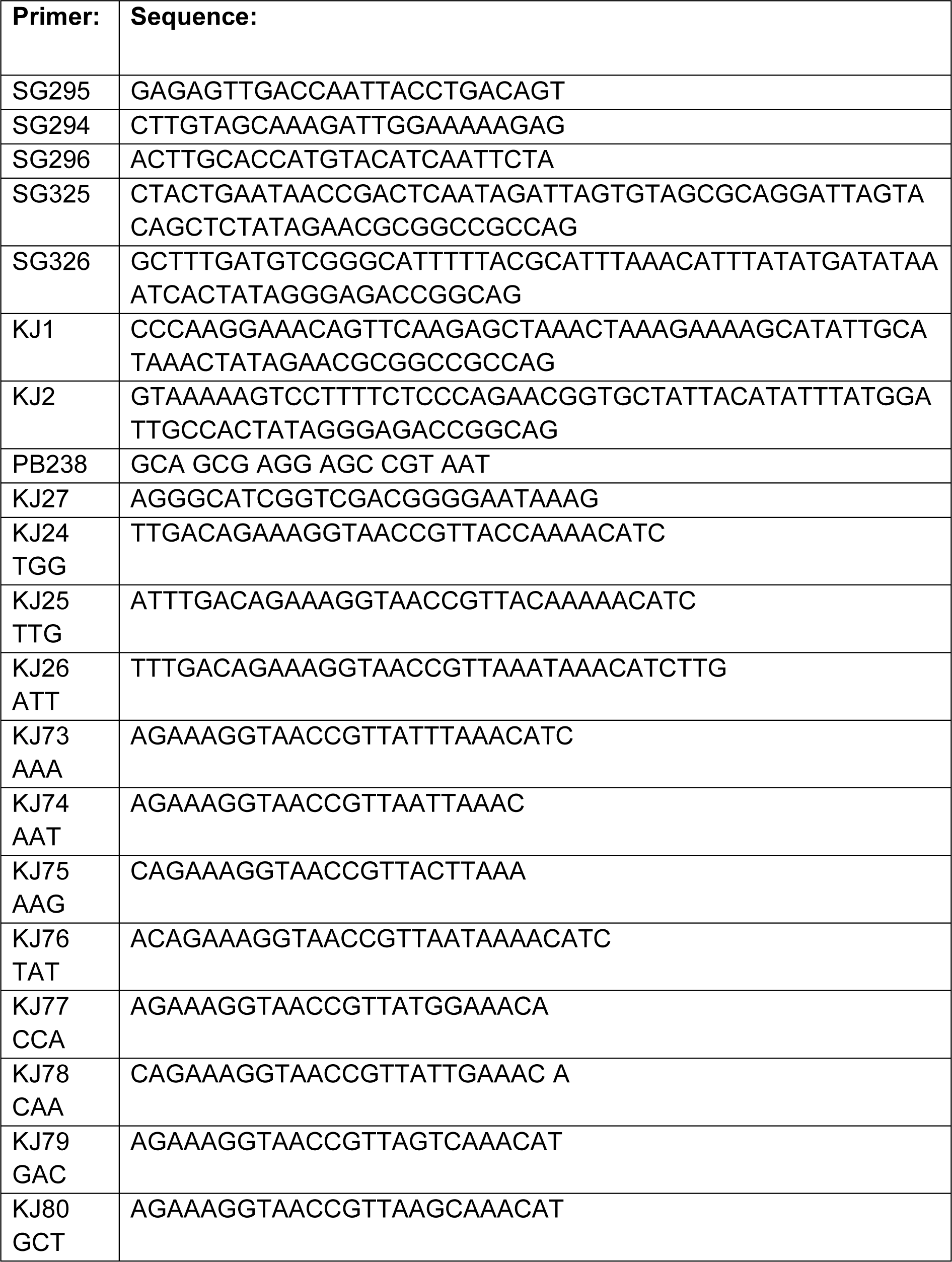
Primers used in this study. This table lists all primers used in this study.

**Table S4.**
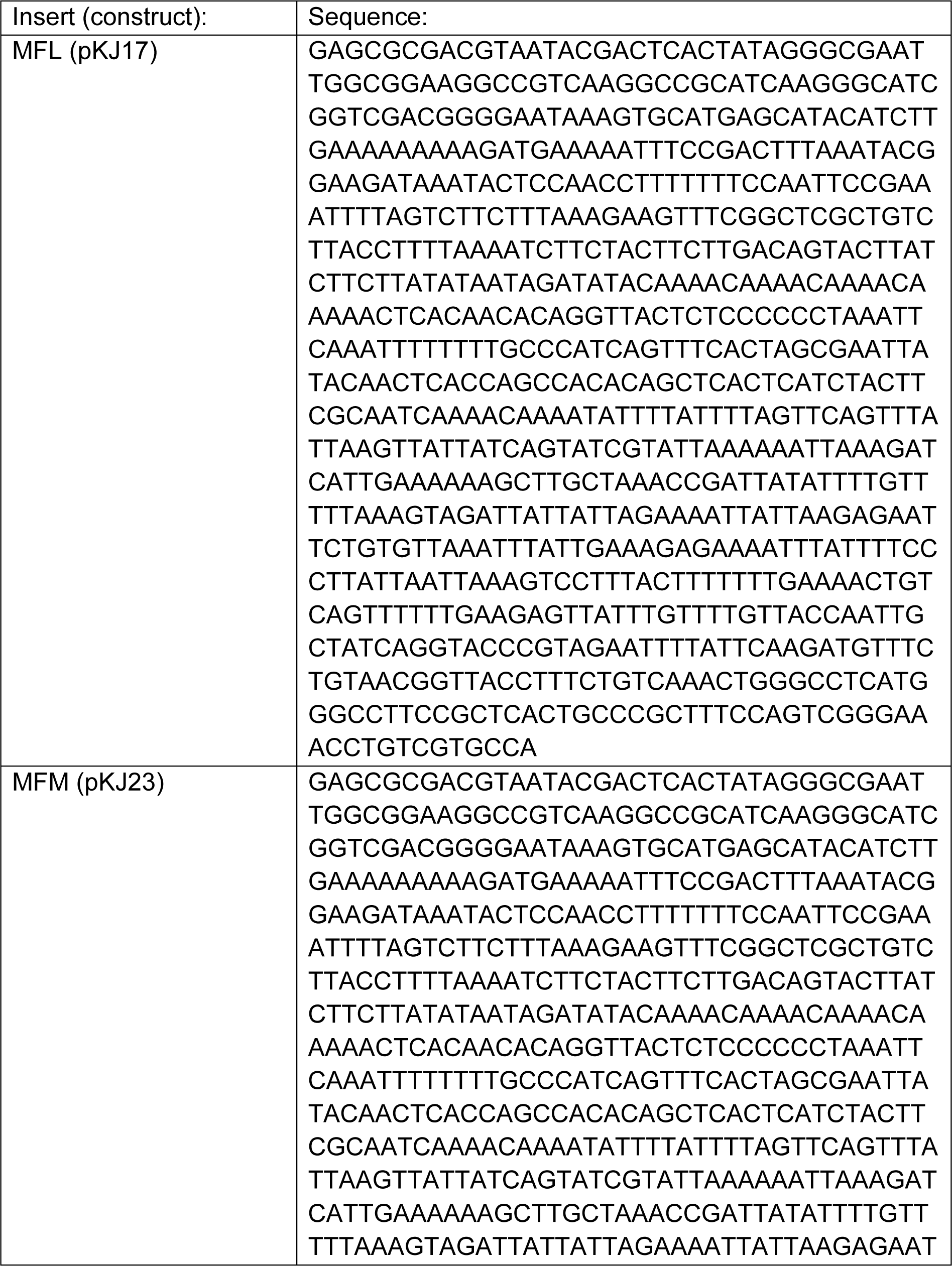
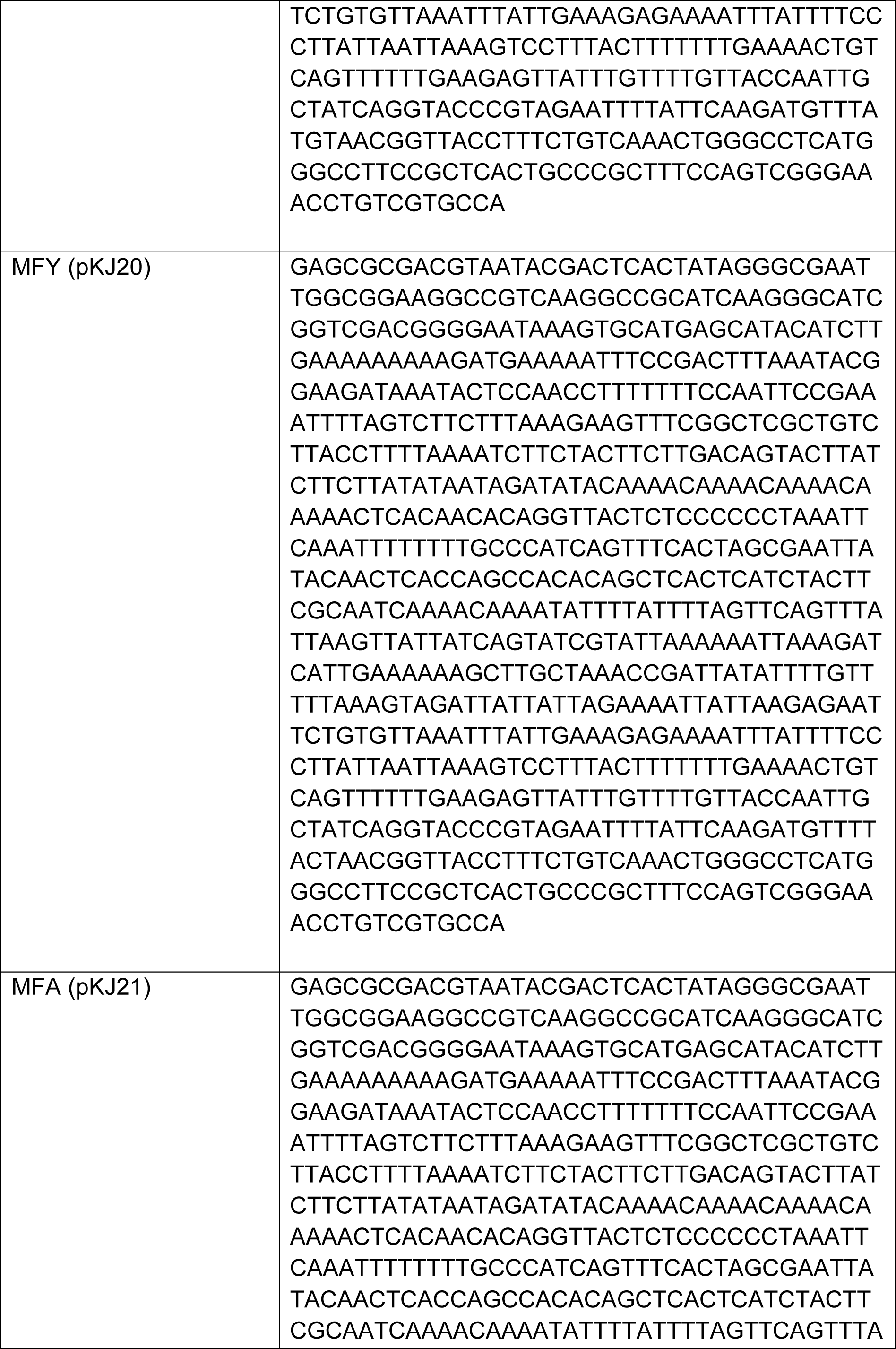
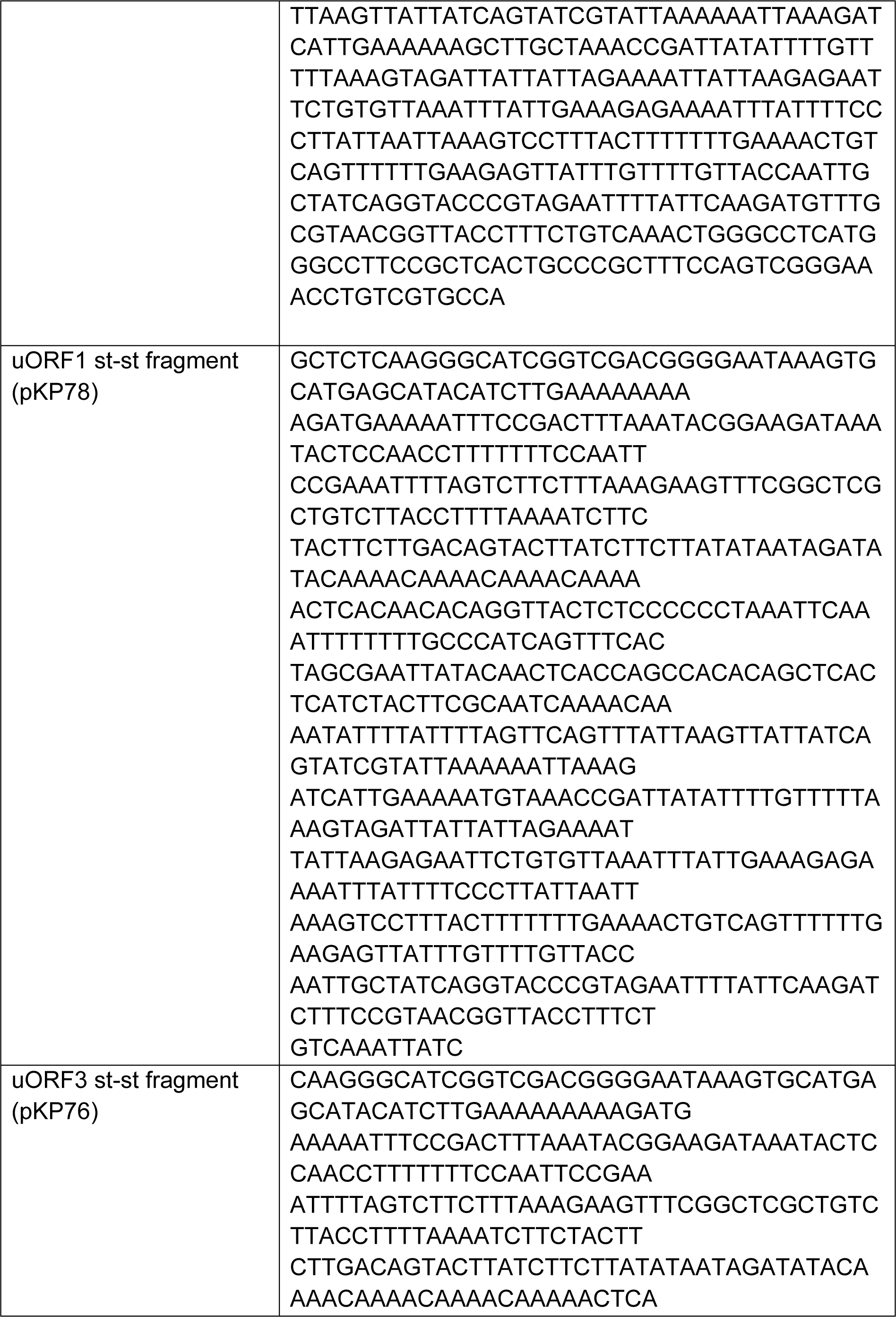
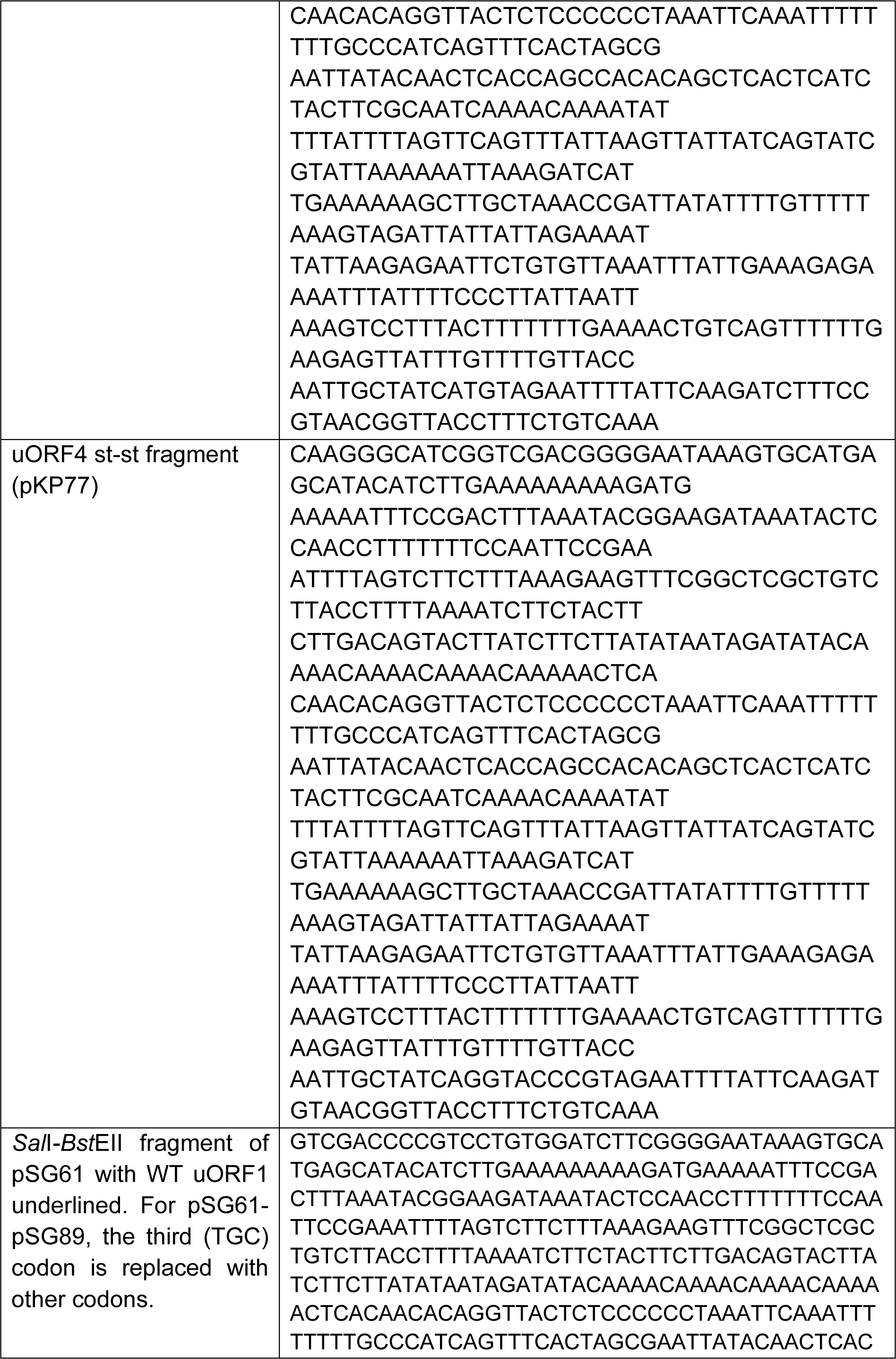
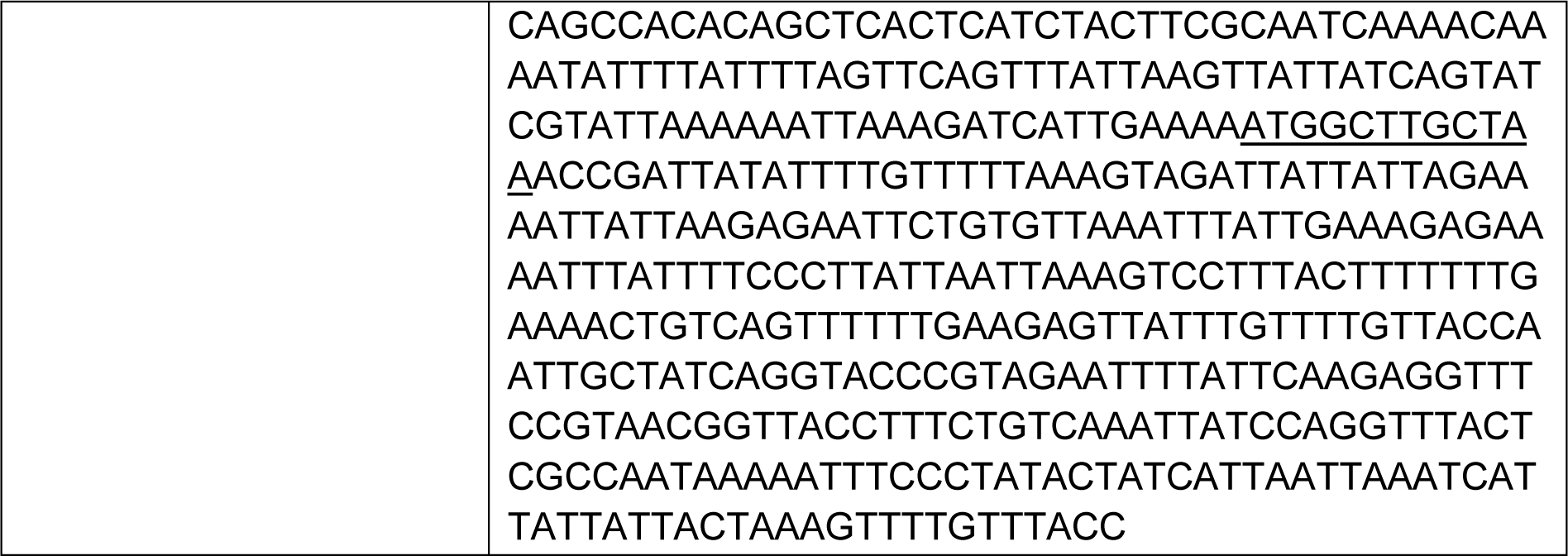
Synthetized DNA inserts used in this study. This table lists the sequences of DNA fragments used for cloning that were synthesized through GeneArt Gene Synthesis (Thermo Fisher Scientific) or by LifeSct LLC. The table is formatted as follows: Insert name (construct it was used for); Construct sequence.

